# Age and axon-specific forms of cortical remyelination by divergent populations of NG2-glia

**DOI:** 10.1101/2020.12.09.414755

**Authors:** Timothy W. Chapman, Genaro E. Olveda, Elizabeth Pereira, Robert A. Hill

## Abstract

Myelin is critical for neural circuit function and its destruction is widespread in neurodegenerative disease and aging. In these conditions, homeostatic repair mechanisms initiate oligodendrocyte replacement by resident progenitor cells called NG2-glia. To investigate the cellular dynamics of this repair we developed a novel demyelination model by combining intravital myelin imaging with a targeted single-cell ablation technique called 2Phatal. Oligodendrocyte 2Phatal activated a stereotyped degeneration cascade which triggered remyelination by local NG2-glia. Remyelination efficiency was dependent on initial myelin patterning and dynamic imaging revealed rapid repair mechanisms resulting in near-seamless transitions between myelin loss and repair. A subset of morphologically complex NG2-glia executed this remyelination, pointing towards unrecognized functional diversity within this population. Age-related demyelination mirrored the degenerative cascade observed with 2Phatal, while remyelination in aging was defective due to failed oligodendrogenesis. Thus, oligodendrocyte 2Phatal revealed cellular diversity within the oligodendrocyte lineage and uncovered novel forms of rapid remyelination.

## INTRODUCTION

Myelination increases neural circuit processing speed and efficiency^1,2^. Many neurodegenerative conditions result from myelin dysfunction. While the myelin sheath is one of the earliest structures to degenerate in aging, a process that is thought to contribute to cognitive decline^3–5^. Functional deficits due to myelin damage are dependent on the degree of myelin loss and the extent to which repair occurs, with late stages of disease often coinciding with failed myelin repair. The cellular mechanisms that allow for the continued refinement and dynamic myelin plasticity, observed in adulthood^6–8^, are thought to be co-opted for myelin repair in these degenerative conditions. Yet, it is not fully known how these cycles of degeneration and repair occur at the cellular level *in vivo*.

Remyelination is primarily carried out by a resident self-renewing population of oligodendrocyte precursor cells, called NG2-glia, which retain the ability to differentiate into myelinating oligodendrocytes throughout life^9,10^. Recent work has revealed dynamic behavior of NG2-glia, including tissue surveillance, targeted response to damage, and participation in remyelination^11–14^. There is some evidence that heterogeneity within the NG2-glia population exists, at least in terms of subsets of cells exhibiting biases towards self-renewal vs oligodendrocyte differentiation under various conditions, brain regions, and ages^15–19^. Importantly, however, the precise cellular dynamics and spatiotemporal responses by NG2-glia to the death of oligodendrocytes and loss of myelin are not known. Moreover, whether or not distinct subsets of NG2-glia participate in oligodendrocyte replacement is yet to be determined.

During development, certain axons exhibit an increased propensity for becoming myelinated, with proper neural circuit function being dependent on this specificity and the myelin patterning along these axons^6,20–22^. Mounting evidence suggests that a combination of neural activity and biophysical properties are key modulators of this decision to myelinate target axons and in which pattern^23–26^. It remains to be determined, however, whether the same axons would also have an increased tendency to be remyelinated following degenerative events. Recent intravital imaging experiments do suggest that both sensory experience^27^ and developmental myelin patterning^28,29^ are important for remyelination success, lending credence to the idea that remyelination proceeds through similar checkpoints used during development.

Here we developed a novel method of focal cortical demyelination using a technique called 2Phatal (<u>2</u>-<u>Ph</u>oton <u>a</u>poptotic <u>t</u>argeted <u>a</u>b<u>l</u>ation)^30^. This approach, in combination with fluorescence and label-free intravital imaging, allowed us to investigate the precise dynamics of oligodendrocyte degeneration, NG2-glia fate in response to myelin loss, and patterns and spatiotemporal dynamics of myelin repair, all in the live mouse cerebral cortex. We show that 2Phatal permits titratable, on demand, cell death induction of individual oligodendrocytes through a remarkably reproducible cell death progression, without compromising adjacent cells or the subsequent repair efforts of NG2-glia. Remyelination timing and success were biased towards axons with more complete myelin coverage. Intriguingly, in some cases, the spatiotemporal dynamics of remyelination resulted in a near seamless transition from sheath degeneration to repair without significant loss of myelin compaction along the affected axonal segment. This remyelination was driven in large part by a subset of NG2-glia which displayed higher degrees of morphological complexity. This morphological identity was predicative of cell behavior, weeks before final fate determination. Finally, in aged mice, spontaneous myelin degeneration looked remarkably similar to oligodendrocyte degeneration that occurred via 2Phatal and cuprizone intoxication, suggesting a common mechanism of cell death in aging and these demyelination models. Consistent with past aging studies, remyelination after 2Phatal was significantly compromised in aged animals, due to the lack of new oligodendrocyte formation in response to myelin loss. Thus, we describe a new model of demyelination and use it to reveal the dynamics of and cell populations involved in remyelination in the adult and aging brain.

## RESULTS

### 2Phatal as a novel model of focal cortical demyelination in the live brain

In order to establish a model of targeted, on-demand, single oligodendrocyte death and demyelination in the cortex of live mice, we implemented a newly developed technique called 2Phatal. This method combines intravital nuclear dye labeling with two-photon mediated photobleaching of single nuclei to induce DNA damage and subsequent cell death via a mechanism resembling apoptosis^30,31^. Topical application of Hoechst 33342 dye to the cortical pial surface during a cranial window surgery provides broad nuclear labeling of cells within the region (Fig. 1a). Combining Hoechst labeling with transgenic reporter mice expressing a membrane tethered EGFP specifically in myelinating oligodendrocytes (*Cnp-*mEGFP)^32,33^, enabled the identification of mature oligodendrocytes and revealed bright and consistent nuclear labeling within this cell population (Fig. 1a). To establish and characterize 2Phatal for oligodendrocyte ablation, we first employed a consistent paradigm for nuclear photobleaching, exposing each nucleus within a single region of interest (ROI) to two photon laser scans (ROI = 8×8μm, 125 scans lasting 3.72s) as described in the methods. This brief focal laser scan resulted in a reproducible decrease in Hoechst fluorescence signal without disrupting cytostructural integrity, as evidenced by no change in the mEGFP of targeted oligodendrocytes (Fig. 1b). Immediately adjacent nuclei and other cell structures maintained normal morphology (Fig. 1b), demonstrating the focality of the subcellular insult. To further investigate if the photobleaching process caused damage to nearby cells, we assessed the immediate activity of microglia following 2Phatal using dual-labeled triple transgenic reporter mice (*Cnp-*mEGFP:*Cx3cr1-* creER:tdTomato) (Fig. 1c). We observed no difference in the immediate chemotactic response by microglia towards targeted or untargeted cells, 20 minutes post photobleaching (Fig. 1d, n = 21 control and 27 2Phatal cells, 4 mice, unpaired t-test). This is in contrast to the immediate chemotactic response by microglia following thermal ablation techniques, due to cell rupture and release of intracellular contents into the extracellular space^30,34^.

**Fig. 1.**
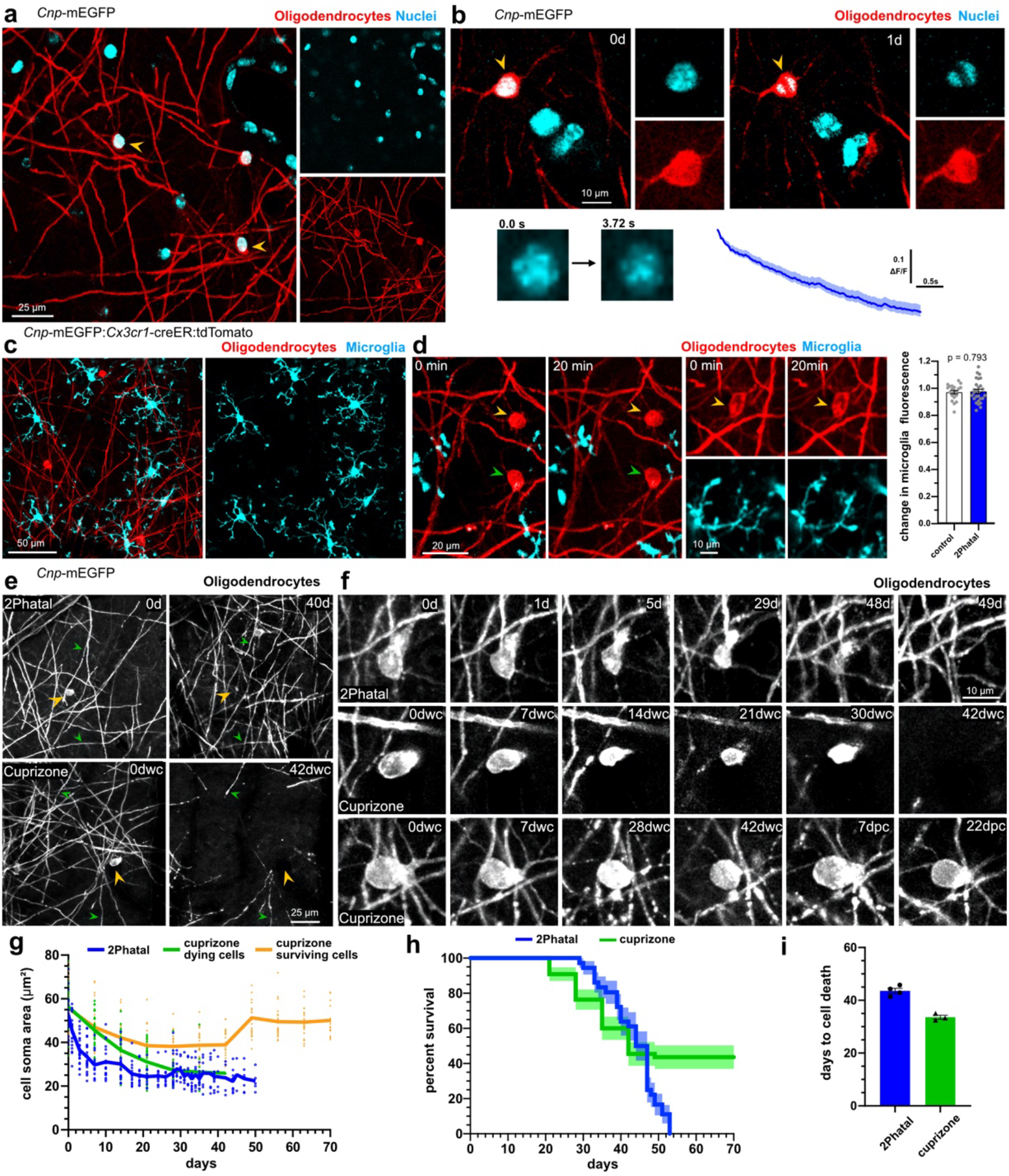
On-demand focal cortical demyelination with 2Phatal. **a)** In vivo image of hoechst nuclear dye labeling (cyan) in the somatosensory cortex of a *Cnp*-mEGFP transgenic mouse with oligodendrocytes labeled via membrane tethered EGFP (arrowheads). **b**) Photobleaching of single oligodendrocyte nuclei (arrowhead, 3.72s bleach) caused disruption of nuclear labeling without causing damage to adjacent cells. Reliable photobleaching is shown by the average fluorescence intensity trace for all targeted cells (n = 82 cells, 6 mice). **c)** *In vivo* images from dual-reporter, triple transgenic mice with oligodendrocytes (red) and microglia (cyan) labeling **d**) Microglia exhibited no immediate chemotactic response towards oligodendrocytes targeted with 2Patal (yellow arrowheads), compared to controls (green arrowheads), demonstrating no disruption of the targeted oligodendrocyte cell membrane during 2Phatal photobleaching. The change in microglia fluorescence intensity was quantified before and 20 minutes after photobleaching in 2Phatal and control cells (n = 21 control cells and 27 2Phatal cells, 3 mice, unpaired t-test). **e**) *In vivo* time lapse images showing oligodendrocyte cell loss (yellow arrowheads) and demyelination in 2Phatal (top) and cuprizone fed (bottom) mice. Green arrowheads denote points of reference for position orientation. **f**) representative timeseries of oligodendrocytes targeted with 2Phatal, oligodendrocytes dying during cuprizone treatment, and oligodendrocytes that survive cuprizone intoxication. 2Phatal was induced on 0d, with final degeneration occurring 7 weeks later (dwc = days with cuprizone; dpc = days post cuprizone). In all 3 cases, soma morphology is disrupted within 7 days of initial insult. Cells that survive cuprizone treatment resume normal morphology after the mice are returned to normal chow. **g**) Changes in oligodendrocyte soma area (μm^2^) of cells targeted with 2Phatal (blue, n = 24 cells, 4 mice), dying cells in mice treated with cuprizone (green, n = 31 cells, 3 mice), and surviving cells in mice treated with cuprizone (orange, n = 24 cells, 3 mice). **h**) survival curve of oligodendrocytes after 2Phatal (blue, n = 36 cells, 4 mice) or in mice treated with cuprizone (green, n = 55 cells, 3 mice) (error represents SEM). **i**) average time to cell death of oligodendrocytes targeted with 2Phatal (n = 4 mice) or cells that died in mice treated with cuprizone (n = 4 mice).

After photobleaching of single oligodendrocytes, longitudinal *in vivo* imaging over several months revealed that oligodendrocyte death proceeded through a predictable cell degeneration process. Following initial condensation of the cell soma over the first 7-10 days, affected oligodendrocytes persisted despite severely disrupted morphology, before finally disappearing, on average, 44 days after 2Phatal photobleaching (n = 36 cells, 4 mice, Fig. 1e-g). Due to the specificity of 2Phatal, only targeted cells degenerated, allowing for the generation of new oligodendrocytes throughout the experiment (Fig. 1e). Interestingly, this protracted timeline for oligodendrocyte cell death was distinct from other cell types, targeted with 2Phatal, such as NG2-glia and neurons, which undergo an apoptotic event within 24 hours after photobleaching^30^.

To determine if a separate, commonly used, model of demyelination proceeded through similar stages of cell death, we directly compared 2Phatal to cuprizone intoxication (Fig. 1e-I, Supplementary Fig. 1). An independent cohort of *Cnp*-mEGFP mice were fed a 0.2% (w/w) cuprizone diet for 6 weeks, followed by 4 weeks of normal chow and imaged weekly to evaluate the dynamics of demyelination and repair (Supplementary Fig. 1a). As expected, mice treated with cuprizone showed widespread oligodendrocyte and myelin loss during the 6 weeks of treatment (Fig. 1e, Supplementary Fig. Fig. 1b). Dying oligodendrocytes in mice treated with cuprizone displayed strikingly similar dynamics to 2Phatal, with cell death occurring only after persistent cell soma condensation, with an average time to cell death of 34 days (n = 31 cells, 3 mice, Fig. 1f-i, Supplementary Fig. 1c-d). Unlike 2Phatal however, which resulted in the death of 100% of the targeted cells (n = 36 cells, 4 mice), oligodendrocyte degeneration was incomplete in the cuprizone model, with 56.4% of the cells dying throughout the imaging period (n = 55 cells, 3 mice). Interestingly, oligodendrocytes that survived the cuprizone treatment exhibited soma condensation, although to a lesser extent compared to the cells that did eventually die (Fig. 1f-g). For surviving oligodendrocytes, the soma condensation was reversible, as soma areas returned to baseline values following the cessation of cuprizone (Fig. 1f-g). The similarities between 2Phatal and cuprizone, in terms of timing and morphological progression during the cell death process, suggest that our observations with 2Phatal are due to an intrinsic response by oligodendrocytes to adverse stimuli, likely mimicking events that occur in other neurodegenerative conditions and aging as outlined below. Thus, 2Phatal of oligodendrocytes is a robust model for titratable, on-demand demyelination, permitting dynamic studies of demyelination and repair, in real time, in the live brain.

### Myelin degeneration proceeds through distinct phases of retraction and decompaction

After analysis of oligodendrocyte death via 2Phatal, we next aimed to investigate how myelin sheath degeneration progressed. Membrane tethered GFP allowed for the visualization of oligodendrocyte proximal processes and tracing of their connection with the myelin sheath. Because of this, we were able to assign a subpopulation of individual sheaths to their cell of origin (Fig. 2a) and perform in depth characterization of single sheath demyelination. Analyses of 183 sheaths from 19 2Phatal targeted oligodendrocytes revealed that 94% of sheaths degenerated prior to the loss of their attached cell soma (Fig. 2a-b). This suggested a temporally linked sequence of oligodendrocyte soma and sheath degeneration. Even with this temporally synchronized process however, the degeneration of all sheaths attached to a single oligodendrocyte occurred over days, indicating the existence of a drawn-out degenerative mechanism, rather than a single catastrophic cell death event (Fig. 2b). Quantitative analysis of myelin sheath length during this process revealed prolonged degeneration, sheath retraction, and formation of characteristic morphological features of damaged myelin (Fig. 2c-f), including sheath thinning and formation of myelin swellings. Sheath retraction was most prominent in the 10 days prior to the loss of each sheath (Fig. 2e, n = 49 sheaths, 4 mice), while thinning and formation of myelin swellings occurred throughout the degenerative cascade.

**Fig. 2.**
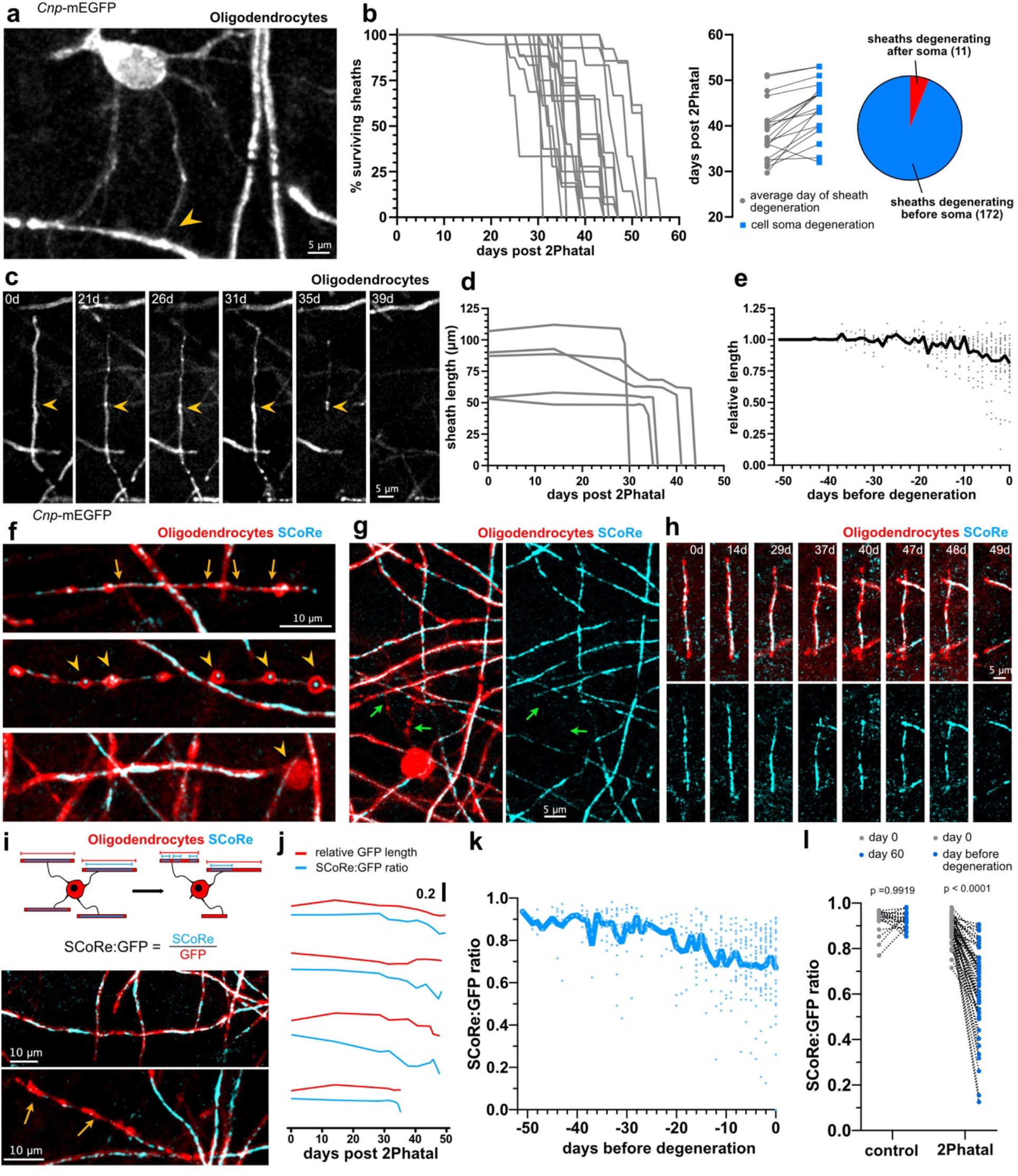
Myelin sheath degeneration occurs via distinct phases of retraction and membrane decompaction. **a)** *In-vivo* imaging of *CNP-*mEGFP reveals oligodendrocyte proximal processes and their connection with the myelin sheath (arrowhead). **b)** Temporal dynamics of sheath degeneration. Each trace represents sheaths produced from a single oligodendrocyte (n = 19 cells, 183 sheaths, 4 mice). Average time to sheath degeneration compared to cell soma loss and breakdown of sheaths degenerating before or after their attached soma for all sheaths analyzed. **c**) In vivo image sequence of a myelin sheath thinning and retracting over weeks after 2Phatal. **d**) Representative length changes of sheaths produced by one 2Phatal-targeted oligodendrocyte. **e**) Relative length changes in all myelin sheaths leading up to the day of their degeneration (grey dots indicate single sheath measurements, n = 49 sheaths, 4 mice). The overlayed black trace represents the mean. **f)**Common myelin sheath pathology observed during 2Phatal induced degeneration, including sheath thinning (top, between arrows), spheroid formation (middle, arrowheads) and myelin debris accumulation (bottom, arrowhead). **g**) *In vivo* image indicating that the SCoRe (myelin compaction) signal is absent from proximal processes (green arrows). **h**) Representative time sequence of a degenerating sheath visualized with fluorescence and SCoRe microscopy. **i)** SCoRe to GFP ratios were calculated to determine SCoRe coverage along myelin sheaths (top). This value was used to distinguish compact sheaths (middle) from uncompacted sheaths (bottom, yellow arrows). **j)** Representative traces of GFP length (red) and SCoRe:GFP ratio (blue) from 4 sheaths. **k)** SCoRe:GFP ratios of all internodes normalized to the day of sheath degeneration (n = 49 sheaths, 4 mice). The overlayed blue trace represents the mean. **l)** SCoRe:GFP ratios of control and degenerating sheaths at day 0 vs day 60 for control or the day before degeneration for 2Phatal (n = 19 control sheaths and n = 49 2Phatal sheaths, 4 mice, paired t-tests with Holm Sidak multiple comparisons).

SCoRe microscopy enables direct label-free visualization of compact myelin, as areas with oligodendrocyte cell membrane that lack membrane compaction, such as at paranodes or along proximal processes, do not generate a SCoRe signal (Fig. 2g)^33,35,36^. Combining this technique with fluorescence microscopy enabled the quantification of changes in compaction of individual sheaths throughout degeneration (Fig. 2h). As membrane compaction is vital to normal myelin function, it was therefore possible to not only detect and quantify changes in morphology, but also gain quantitative and spatiotemporal insight into deterioration of myelin sheath function. Changes in SCoRe were reported as a ratio of total SCoRe length to mEGFP length, in order to more easily reflect the extent of compaction along the entire sheath (Fig. 2i). While the timing of decompaction varied between sheaths measured, the average SCoRe:GFP ratio decreased steadily over the course of degeneration (Fig. 2j-k). Loss of compaction was most severe immediately prior to sheath loss, with no loss of compaction observed in control non-targeted sheaths (Fig. 2l, n = 19 control sheaths and 49 2Phatal sheaths, 4 mice, paired t-tests with Holm Sidak multiple comparisons correction). Interestingly, decompaction was also observed up to 20 days prior to sheath degeneration, suggesting that severely disrupted, non-compact myelin can persist without being removed from the axon for weeks (Fig. 2j-k). These data show that oligodendrocyte death results in protracted myelin sheath deterioration, characterized by gradual decompaction, sheath thinning, and complete demyelination.

### Focal NG2-glia response to oligodendrocyte death and demyelination

As NG2-glia are responsible for replacing lost myelin through oligodendrocyte differentiation, it was critical to investigate how oligodendrocyte death and demyelination impacted their behavior. To observe and quantify this response, we employed a new transgenic mouse line capable of discriminating NG2-glia from myelinating oligodendrocytes. We generated triple transgenic, dual-reporter mice with mEGFP in myelinating oligodendrocytes and tamoxifen inducible tdTomato in NG2-glia and their progeny, *Cnp*-mEGFP*:Cspg4*-creER:tdTomato (Fig. 3a-c, Supplementary Fig. 2). Mice were injected with tamoxifen 2 days prior to surgery, ensuring sufficient time for reporter expression, while minimizing double labeled myelinating oligodendrocytes at the start of the experiment. Oligodendrocyte differentiation from NG2-glia was identified by the appearance of mEGFP expression, resulting in double labeled (tdTomato+mEGFP+) myelinating oligodendrocytes (Fig. 3c, Supplementary Fig. 2).

**Fig. 3.**
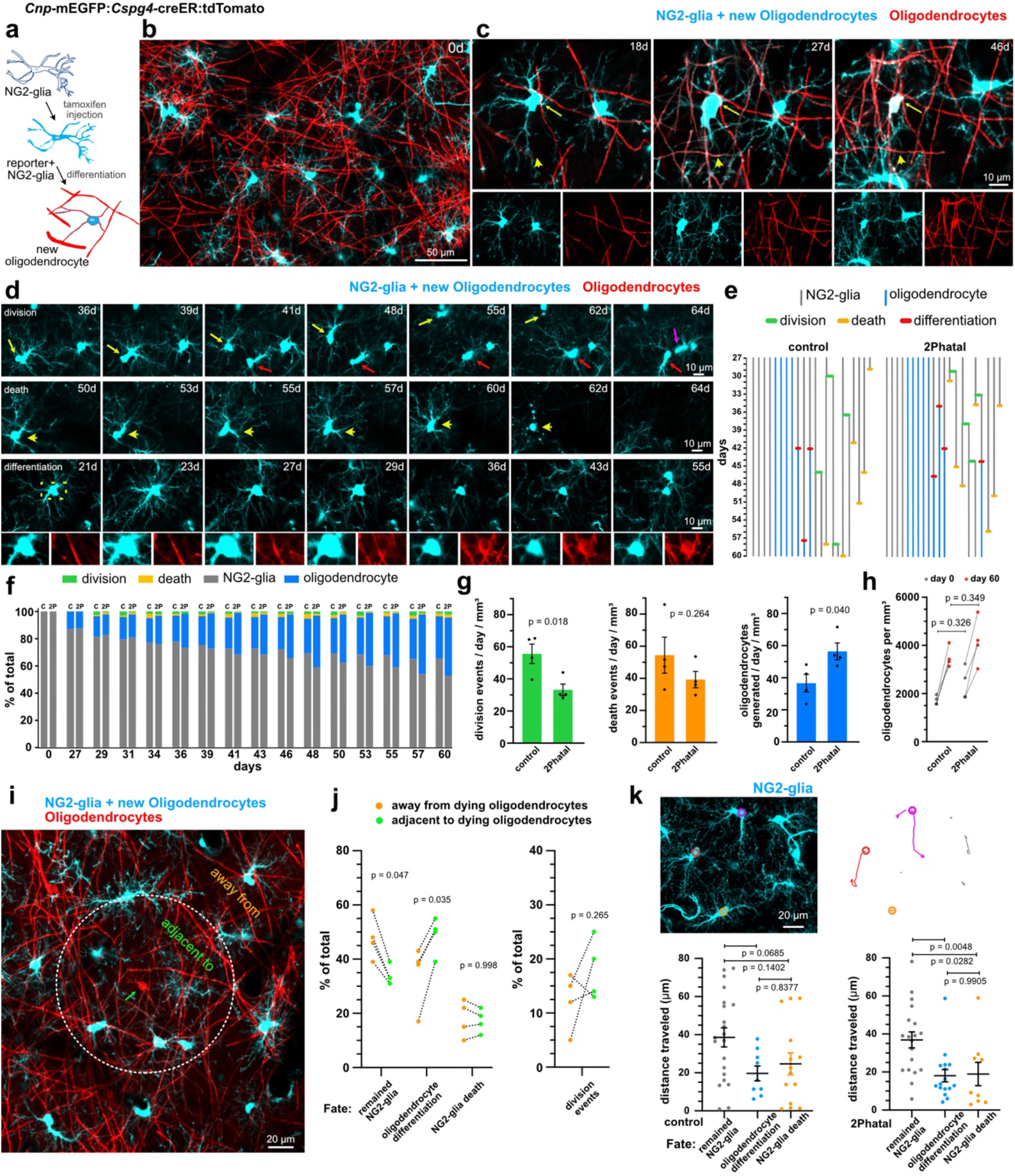
Divergent fates of single NG2-glia after oligodendrocyte 2Phatal. **a)** Dual color fate mapping of NG2-glia in *Cnp-*mEGFP:*Cspg4-*creER:tdTomato mice with an inducible cre reporter. **b)** In vivo image of NG2-glia (cyan) and myelinating oligodendrocytes (red) immediately following cre recombination. **c)** NG2-glia differentiation into a dual-labeled myelinating oligodendrocyte (arrows). New myelin sheaths generated during differentiation are dual labeled (arrowheads). **d**) In vivo time-lapse images showing outcomes for NG2-glia fate including cell division (top) cell death (middle) and oligodendrocyte differentiation (bottom, boxed area shows the *Cnp*-mEGFP channel indicating the turning on of mEGFP as the cell differentiates). **e**) NG2-glia lineage diagrams indicating division events (green), apoptosis (yellow), and oligodendrocyte differentiation (red) in control and 2Phatal mice. **f**) NG2-glia fate in control (c) and 2Phatal (2P) mice, as a percentage of all cells. (control, n = 182 cells, 4 mice; 2Phatal, n = 256 cells, 4 mice). **g**) NG2-glia division events per day (green), apoptotic events per day (orange), and differentiation events per day (blue) in control mice compared to 2Phatal mice (n = 4 mice, unpaired t-tests). **h)** Oligodendrocyte density in control and 2Phatal mice at day 0 and day 60 (n = 4 mice, two-way ANOVA, Sidaks multiple comparison’s test). **i**) Analysis of NG2-glia adjacent to or away from oligodendrocytes targeted by 2Phatal. Adjacent cells were defined as those with somas on or within a 150μm diameter circle centered around each targeted cell (green arrow). **j**) The proportion of all cells that remained NG2-glia, differentiated, or died in both regions (n = 4 mice, two-way ANOVA Sidaks multiple comparison’s test). Division events between these populations was also quantified (n = 4 mice, paired t test). **k**) Example tracks from semi-automated measurements of NG2-glia migration in control and 2Phatal mice. Analysis of total distance traveled relative to cell fate for each group. Dots represent a single cell and horizontal lines indicate the mean (control n = 48 cells, 2Phatal n = 45 cells, error bars are SEM, one-way ANOVAs with Tukey correction for multiple comparisons).

NG2-glia fate was analyzed during peak demyelination and remyelination (days 28-60 after 2Phatal) in animals subjected to oligodendrocyte 2Phatal and control animals (Fig. 3d). Documenting division, differentiation, and death events enabled the generation of individualized lineage diagrams of all NG2-glia within each imaging location (n = 182 cells, 4 control mice and 256 cells, 4 2Phatal mice, Fig. 3e). Overall, we found a significant increase in the generation of new oligodendrocytes in the 2Phatal mice, accompanied by a decrease in imaged division events (Fig. 3f-g, n = 4 mice, unpaired t-test), however there were no significant differences in the total density of oligodendrocytes between control and 2Phatal mice at day 0 and day 60, indicating that the excess oligodendrocyte production was able to compensate for the loss of oligodendrocytes due to 2Phatal (Fig. 3h, n = 4 mice, two-way ANOVA, Sidak’s multiple comparison’s test).

To more precisely determine the specificity for NG2-glia fate decisions in response to demyelination, we next analyzed the fate of NG2-glia located either within a 150μm diameter circle of targeted oligodendrocytes or outside of this predetermined territory (Fig. 3i-j). The average fate outcomes were determined for each group of cells, using the same criteria described above. NG2-glia adjacent to dying oligodendrocytes were biased towards oligodendrocyte differentiation while NG2-glia away from dying cells were more likely to remain NG2-glia (Fig. 3i-j, n= 4 mice, two-way ANOVA, Sidak’s multiple comparison’s test). These data demonstrate the focality of the response and provide further evidence that microregional cues modulate the fate outcome of NG2-glia in a targeted and precise fashion.

In addition to fate decisions, NG2-glia are also capable of cell migration in the adult brain, particularly in the context of demyelination and focal injury. In order to characterize this behavior within our system, we performed semi-automated analysis of NG2-glia migration in both the control and 2Phatal conditions. Overall, there were no significant differences in the average total distance traveled or displacement from the original location between NG2-glia in control and 2Phatal mice (Supplementary Fig. 3, n = 48 NG2-glia, 4 control mice and 45 NG2-glia, 4 2Phatal mice, unpaired t-test). By connecting NG2-glia fate with migratory behavior, we did discover a difference in total distance traveled between cells that remained NG2-glia vs those that either differentiated into oligodendrocytes or died in the 2Phatal condition, with the latter two groups exhibiting less migratory behavior over the 32 days of analysis (Fig 3k, one-way ANOVA, Tukey’s multiple comparison’s test).

### Morphological complexity predicts NG2-glia fate during remyelination

During our analysis of NG2-glia response to demyelination, it became apparent that there were varying degrees of morphological complexity within the NG2-glia population (Fig. 4). *In silico* tracing paired with Sholl analyses confirmed this initial observation (Fig. 4a-b). Highly branched cells often had 200% more intersections than cells with less complex morphology; in rarer cases, this difference was as much 600% (Fig. 4b,d). Importantly, none of the cells that were analyzed expressed *Cnp*-mEGFP or were associated with the vasculature, indicating that they were not myelinating oligodendrocytes or perivascular mural cells.

**Fig. 4.**
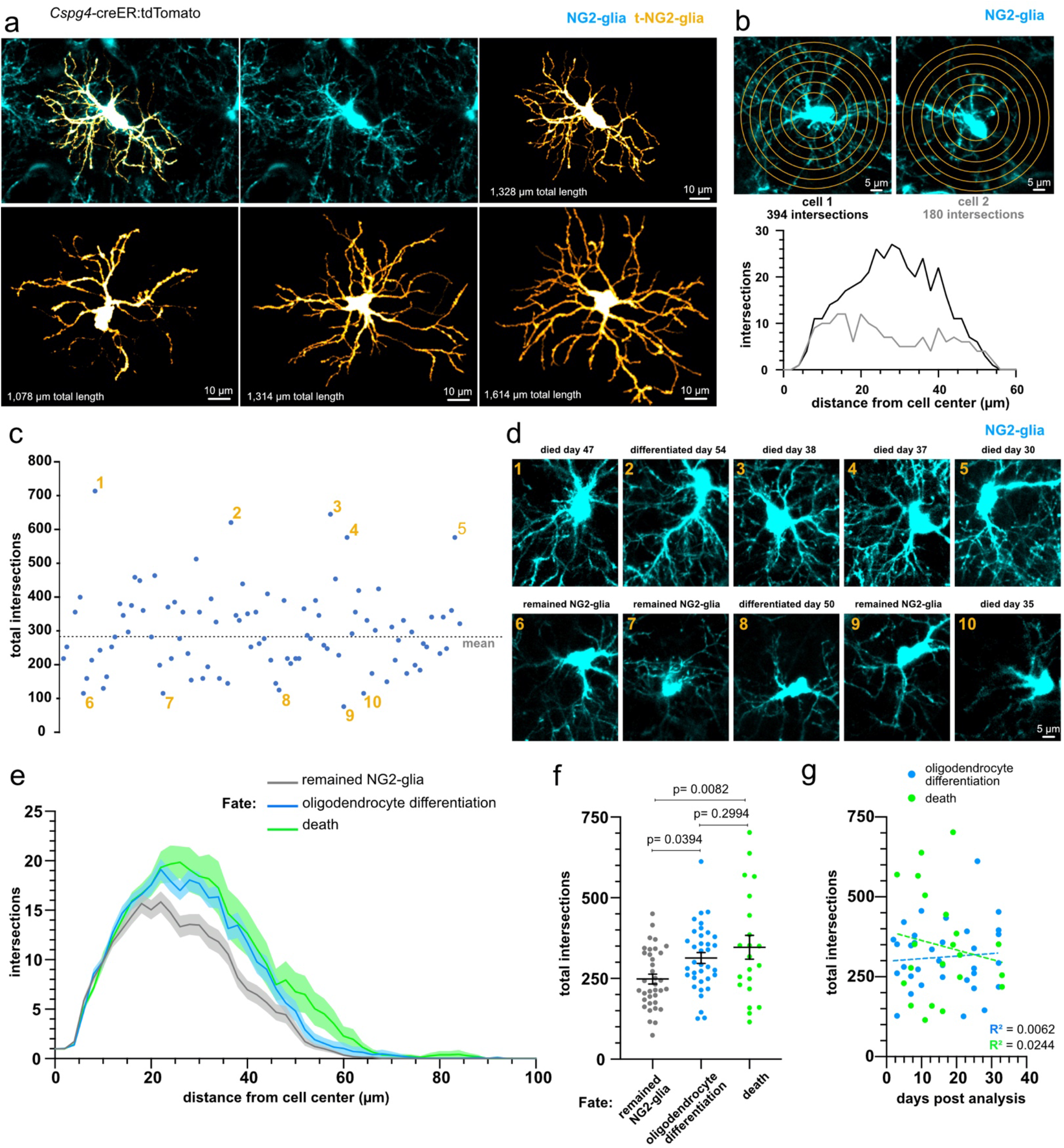
Myelin repair is carried out by a subset of morphologically complex NG2-glia. **a**) Semi-automated reconstructions of NG2-glia with distinct process morphology. Total length represents the summation of process lengths from each cell. **b**) Sholl analyses of two representative NG2-glia with the total number of cell process intersections plotted relative to the cell center. **c**) The distribution of total intersections of NG2-glia analyzed using Sholl analysis. Each dot represents a single cell. The dotted line represents the mean (n = 100 cells, 4 mice). The 5 cells with the most (1-5) and least (6-10) total intersections are labeled. **d**) Images showing the morphology of the 10 NG2-glia identified in **c**, paired with the fate of each of those cells. **e**) Average Sholl analysis plot of NG2-glia separated by fate (error bars are SEM). **f**) Grouped data showing the total intersections of NG2-glia that remained NG2-glia (grey), differentiated, and died (n = 93 cells, 4 mice, one-way ANOVA with Holm-Sidak correction for multiple comparisons, error bars are SEM). Each dot represents a single cell, and the horizontal line represents the mean. **g**) Linear regression analyses revealed no correlation between total intersections and the time from Sholl analysis to final fate outcome (n = 34 differentiating cells and n = 21 dying cells).

Using the previously determined fates of single NG2-glia described in Figure 3, we next aimed to identify any differences in fate outcomes between cells with simple and complex morphology (Fig 4c-g). Intriguingly, initial analysis of cells at the extremes of morphological complexity pointed towards a connection between complexity and eventual fate. For example, 4 of the 5 most branched NG2-glia died over the course of the experiment, while 3 of the 5 simplest cells remained NG2-glia (Fig. 4c-d). Further analysis revealed that this trend was not limited to the extremes (Fig. 4e-f). Binning all tracked cells by outcome showed that NG2-glia which eventually differentiated into oligodendrocytes or died via apoptosis were more likely to have increased process complexity. Cells with fewer total processes were more likely to remain NG2-glia (Fig. 4f, n = 93 cells, 4 mice, one-way ANOVA, Holm-Sidak multiple comparison’s test). In contrast to these associations, we did not observe any differences in complexity between cells that did or did not divide during the imaging period (non-dividing cells = 303 ± 13.5 intersections vs dividing cells = 272 ± 30.5 intersections, p = 0.337, unpaired t test). While it is generally recognized that NG2-glia undergo changes in morphological complexity during oligodendrocyte differentiation, we did not find a correlation between process complexity and time to fate outcome post analysis (Fig. 4g). In fact, the most branched cell we recorded did not undergo apoptosis until 19 days after Sholl analysis (Fig. 4g). Taken together, this suggests that differences in process arborization complexity are not a transient effect occurring while these cells are actively differentiating or undergoing apoptosis, but rather may point to morphologically distinct subpopulations of NG2-glia with differential fate tendencies.

### Remyelination success and timing is dependent on the preexisting myelin pattern

Unlike other models of demyelination, such as cuprizone intoxication, 2Phatal does not influence the ability of NG2 cells to naturally (and rapidly) repair lost myelin through differentiation. This enabled us to use *Cnp*-mEGFP:*Cspg4-*creER:tdTomato dual reporter mice to determine the precise dynamics and patterns of remyelination during oligodendrocyte death and demyelination. As before, inducing cre recombination at the start of the experiment ensured myelin present at the time of 2Phatal ablation was singly labeled with mEGFP, while any new myelin was dual labeled with mEGFP and tdTomato (Fig. 5a). First, we examined the full extent of demyelination and repair within each imaging location. A total of 92 sheaths, from 4 mice, were randomly selected on day 0 and evaluated for stability, degeneration without repair, or degeneration with remyelination (Fig. 5b). This population of sheaths contained sheaths attached to targeted and non-targeted cells. On average, 48.9 ± 4.2% of sheaths identified remained stable over the course of the experiment, 44.1 ± 5.4% degenerated and were repaired, while 7 ± 2.7% fully degenerated without being remyelinated (Fig. 5c). Focusing on sheaths attached to 2Phatal targeted cells revealed that 79.9% of all sheaths that degenerated were remyelinated at some point during our imaging (Fig. 5d).

**Fig. 5.**
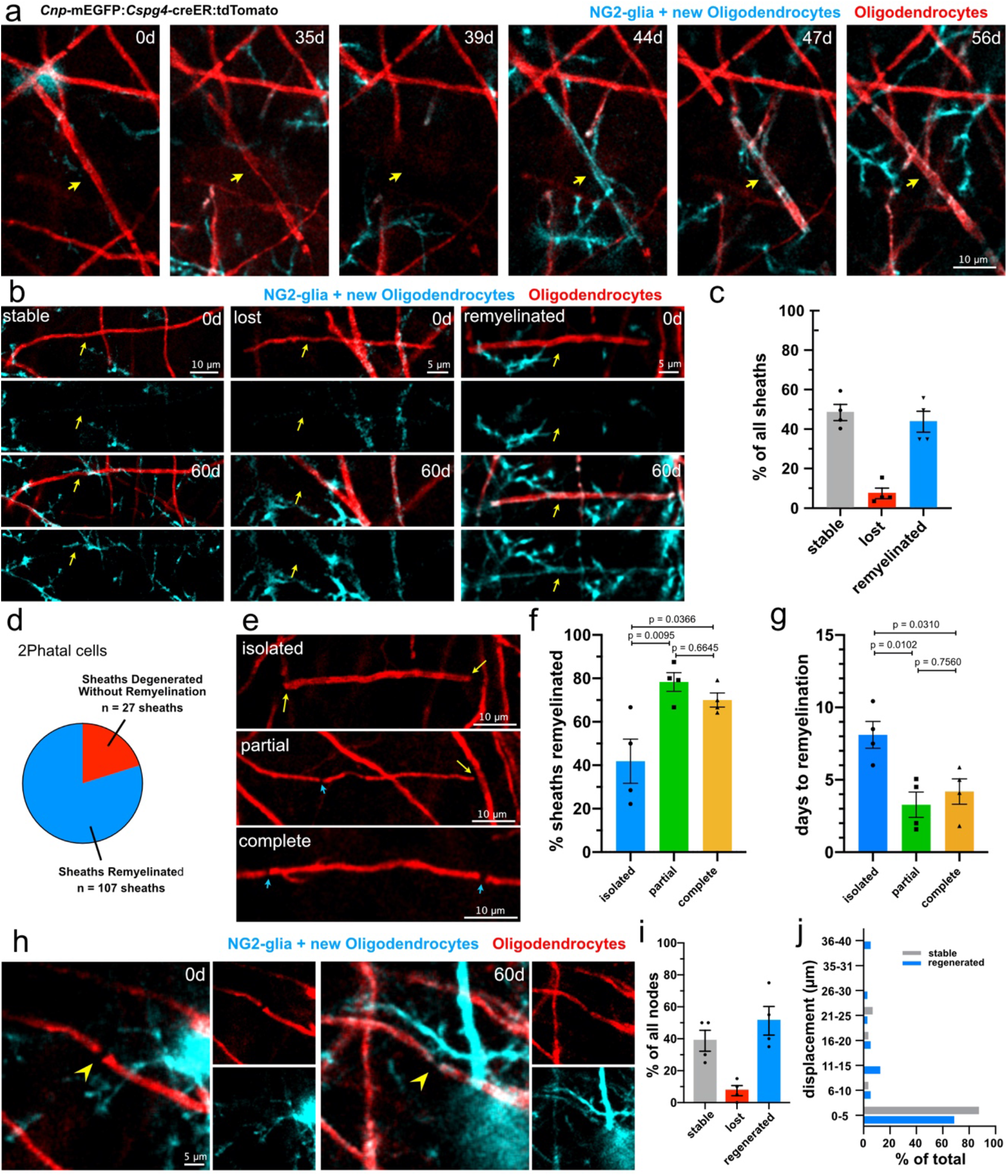
Myelin patterns impact the success and spatiotemporal dynamics of remyelination. **a**) In vivo time series showing degeneration of a single labeled mEGFP+ myelin sheath (arrow) followed by remyelination by a new, dual-labeled mEGFP+/tdTomato+ sheath from a newly differentiated oligodendrocyte. **b**) Representative images of sheaths which remained stable (left), were lost without repair (middle), and degenerated and were remyelinated (right). **c**) Proportion of sheath outcomes for randomly selected sheaths within the cranial window. (n = 92 sheaths, 4 mice, error bars are SEM). **d)** The proportion of sheaths attached to 2Phatal cells that were remyelinated **e**) Representative images showing the 3 possible myelin patterns. Isolated sheaths (top) had no adjacent sheaths (yellow arrow), partial sheaths (middle) had an adjacent sheath on one side (blue arrows) and no adjacent sheath on the other (yellow arrow), and complete sheaths (bottom) had two adjacent sheaths (blue arrows). **f-g**) Remyelination efficiency and time to remyelination were quantified based on the myelin patterning of each sheath. Sheaths with partial and complete patterns were more likely to be remyelinated than isolated sheaths (134 sheaths from n = 4 mice, one-way ANOVA with Turkey correction for multiple comparisons). Remyelination was slower in isolated sheaths compared to the other patterns (107 sheaths from n = 4 mice, one-way ANOVA with Turkey correction for multiple comparisons). Dots represent the average from a mouse. **h**) In vivo images of a node of Ranvier present at day 0 between mEGFP only labeled sheaths, which reformed at its original location between two mEGFP and tdTomato double labeld sheaths, as seen at day 60. **i**) Nodes from 2Phatal mice were randomly selected on day 0, blind to the cell of origin, and evaluated on day 60, to determine if they were stable (mEGFP+ only) were reformed through remyelination (mEGFP+tdTomato+) or degenerated (80 nodes from n = 4 mice) j) Proportion of nodes that were displaced between days 0 and 60 for both nodes that remained stable (n = 31) and nodes that were regenerated following remyelination (n = 41). All error bars are SEM.

In the cortex, myelin is often not uniformly distributed along stretches of axons; thus, we next determined how myelin patterning influenced both remyelination efficiency and the time between degeneration and repair (Fig. 5e-g). Patterning was defined by the number of sheaths adjacent to the sheath of interest; no adjacent sheaths was termed isolated, while 1 and 2 adjacent sheaths were deemed partial and complete respectively (Fig. 5e). Analysis of sheaths, connected to 2Phatal targeted oligodendrocytes, revealed a bias towards remyelination of partial and complete sheaths. Only 42% of isolated sheaths were successfully remyelinated, significantly less than both partial (78%) and complete (70%) patterns (Fig. 5f, one-way ANOVA Tukey’s multiple comparison’s test). In addition to there being a differential outcome in terms of remyelination success, we also detected a difference in the time between sheath degeneration and repair. Isolated sheaths that did get remyelinated, took, on average, 8.5 days to become remyelinated, compared to 3.8 days for sheaths with 1 neighbor and 4.8 days for sheaths with 2 neighbors (Fig. 5g). Of all the sheaths analyzed for remyelination 99.1% (106/107) were dually labeled with mEGFP and tdTomato, indicating the remyelination was carried by newly differentiated cells, not adjacent mature oligodendrocytes (Supplementary Fig. 4d). We only encountered a single remyelinating sheath that was not dually labeled. However due to the emergence of a mEGFP positive, tdTomato negative oligodendrocyte within the territory of that sheath, it seems unlikely that this remyelination was due to the generation of a new sheath from a mature oligodendrocyte present at the start of the experiment, but instead was the result of incomplete cre recombination (Supplementary Fig. 4).

In addition to remyelination of single sheaths, we next determined if nodes of Ranvier were re-established at the same locations prior to degeneration. Nodes were randomly selected at day 0 and evaluated for stability, degeneration, or degeneration and re-establishment. Overall, 38.8% of the nodes were stable, 7.5% never reformed, and 53.8% were re-established during the imaging period (Fig. 5h-i, 80 nodes from n = 4 mice). Nodes that were stable across the imaging period tended to remain stationary, with 87.1% moving less than 5μm (Fig. 5j, 31 nodes, 4 mice). Although less resilient than stable nodes, 68.3% of regenerated nodes re-formed within 5 μm of their original position (Fig. 5j, 43 nodes, 4 mice).

### Rapid restoration of myelin compaction during remyelination

Myelin compaction is necessary for proper saltatory conduction. Because of this, understanding the timing between the emergence of a new oligodendrocyte process and when it transforms into a compact myelin sheath is critical for identifying when functional myelination (and remyelination) has occurred. Using combined time-lapse fluorescence and SCoRe microscopy enabled us to assess, not only the dynamics of the formation of remyelinating sheaths, but also provided insight in the precise timing of the sheath compaction. In a majority of sheaths that did get remyelinated (76.6%, n = 82/107 sheaths, 4 mice) we observed a stereotypical pattern characterized by the loss of the damaged sheath, a period spent in an unmyelinated state, followed by initiation of myelin replacement by the generation of a new oligodendrocyte and myelin sheath (Fig. 6a,f). During this process, we observed a delay in visible SCoRe signal arising from the forming fluorescently tagged sheath (Fig. 6a). Measurement of the same metric that we established during sheath degeneration, the SCoRe:GFP ratio (Fig. 2), revealed that sheath compaction could be delayed by as much as 3 days relative to the start of axon ensheathment, as seen by fluorescence microscopy (Fig. 6d; n = 19 sheaths, 3 mice).

**Fig. 6.**
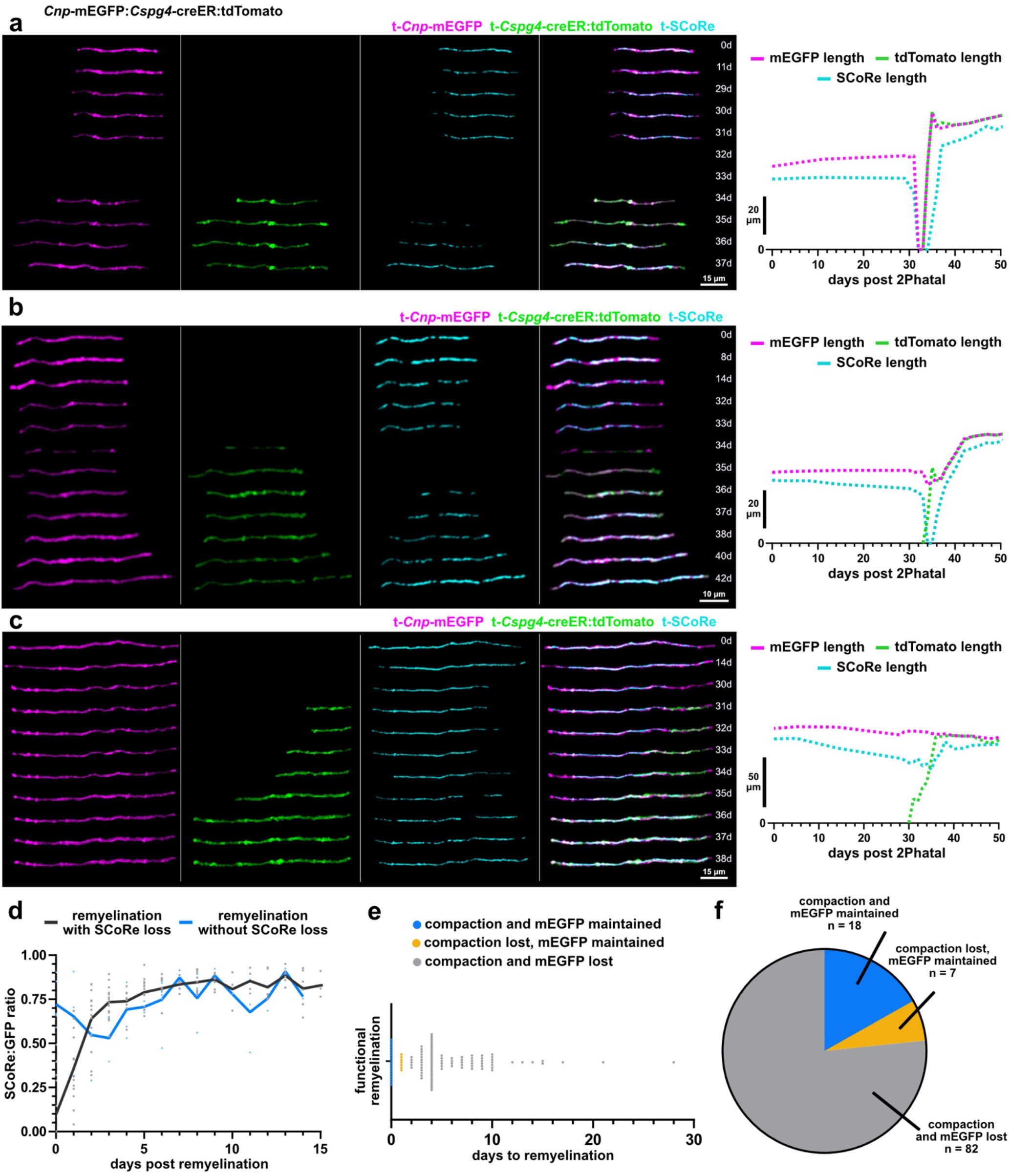
Distinct forms of rapid myelin sheath repair and membrane compaction. **a**) In vivo time lapse images showing a reconstructed myelin sheath that degenerates and is replaced by a new mEGFP+tdTomato+ double labeled sheath. In this sheath there is a two-day window where myelin is absent from the axon. The lengths of mEGFP, tdTomato, and SCoRe at each day are graphed to the right. **b**) A separate example of an additional form of remyelination where unlike the example in (**a**) there is no gap in time where mEGFP is completely absent from the axon suggesting that the emergence of the new sheath occurs simultaneously with the loss of the original. The SCoRe signal is lost, however. **c**) A third example of remyelination that proceeds with a seamless transition between the degenerating and remyelinating sheath. There is no observed gap in the mEGFP signal or the SCoRe signal indicating that compaction is mostly maintained through the repair process. **d**) Data showing the dynamics of SCoRe coverage, of new remyelinating sheaths, following the complete loss of SCoRe signal (black, n = 19 sheaths, 3 mice) or during a seamless transition event (blue, n = 7 sheaths, 2 mice). **e-f**) Time to the emergence of SCoRe signal in remyelinating sheaths, termed “functional remyelination”, grouped by the observed repair pattern described in (**a-c**) and showing the proportion of each type of event (n = 107 sheaths, 4 mice).

In most cases myelin repair was initiated after the complete degeneration of damaged sheaths, however remyelination was also occasionally observed starting before the complete removal of the degenerating sheath (Fig. 6b-c). This process often occurred following loss of SCoRe in the damaged sheath, but before the full removal of myelin debris (Fig. 6b,f; 6.5% n = 7/107, 4 mice). Although compaction was also delayed in these cases, as before, initiating repair in this manner reduced the time between SCoRe loss and recovery (Fig. 6e). We did not detect any abnormalities in sheath morphology that would indicate that the presence of myelin debris impacted normal sheath maturation (Fig. 6b).

Surprisingly, we also detected a form of remyelination in which the SCoRe signal was never lost along the length of the degenerating sheath (Fig. 6c,f; 16.8% n = 18/107, 4 mice). Instead, the newly forming sheath invaded the territory of the damaged sheath, gradually replacing it without any noticeable loss of mEGFP signal and minimal loss of SCoRe (Fig. 6c, Supplementary Fig. 5). We did not observe any breaks in mEGFP separating the remyelinating and preexisting sheath, meaning this form of remyelination could only be detected using a dual reporter system such as ours. The lack of a space between the degenerating and remyelinating sheath suggested that the remyelinating process abuts immediately adjacent to the degenerating sheath, producing an astonishingly rapid form of remyelination, with the potential for minimal functional impact to the underlying axon.

### Disruption of myelin maintenance and repair in aging

Myelin loss is one of the earliest degenerative processes in the aging brain^5^. However, we know very little about the cellular dynamics of this degeneration or the mechanisms inhibiting repair. Time-lapse *in vivo* imaging in aged (22-24 months) *Cnp-*mEGFP mice revealed protracted oligodendrocyte and myelin sheath loss that closely resembled the temporal dynamics and morphological degenerative cascade observed during 2Phatal demyelination (Fig. 7a, Supplementary Fig. 6). This included oligodendrocyte soma shrinkage, myelin sheath thinning, formation of myelin swellings/balloons, and loss of myelin compaction (Fig 7a-e, Supplementary Fig. 6). While stochastic oligodendrocyte death was observed (Fig. 7a-b), overall, aged mice did not show significant changes in oligodendrocyte density over 60 days of imaging (Fig. 7d, n = 3 mice, paired t-test). In contrast, we did observe a significant decrease in myelin sheath density over the same period (Fig. 7e, n = 3 mice, paired t-test) reflecting the gradual loss of single sheaths over time (Fig. 7c). The percent change in oligodendrocyte and myelin sheath density over 60 days were significantly different when comparing 2-month-old with aged animals, as would be expected based on the ongoing generation of new oligodendrocytes in young adult animals (Fig. 7d-e, n = 4 young and 3 aged mice, unpaired t-tests). Moreover, unlike young animals, we rarely observed the generation of a new oligodendrocyte in response to oligodendrocyte loss or myelin degeneration. A single case where this did happen, however, revealed remarkable reestablishment of the myelin pattern (Fig. 7a,g). However, in most cases we observed loss of myelin sheaths and no evidence of new oligodendrocyte formation or remyelination by adjacent mature oligodendrocytes (Fig. 7g).

**Fig. 7.**
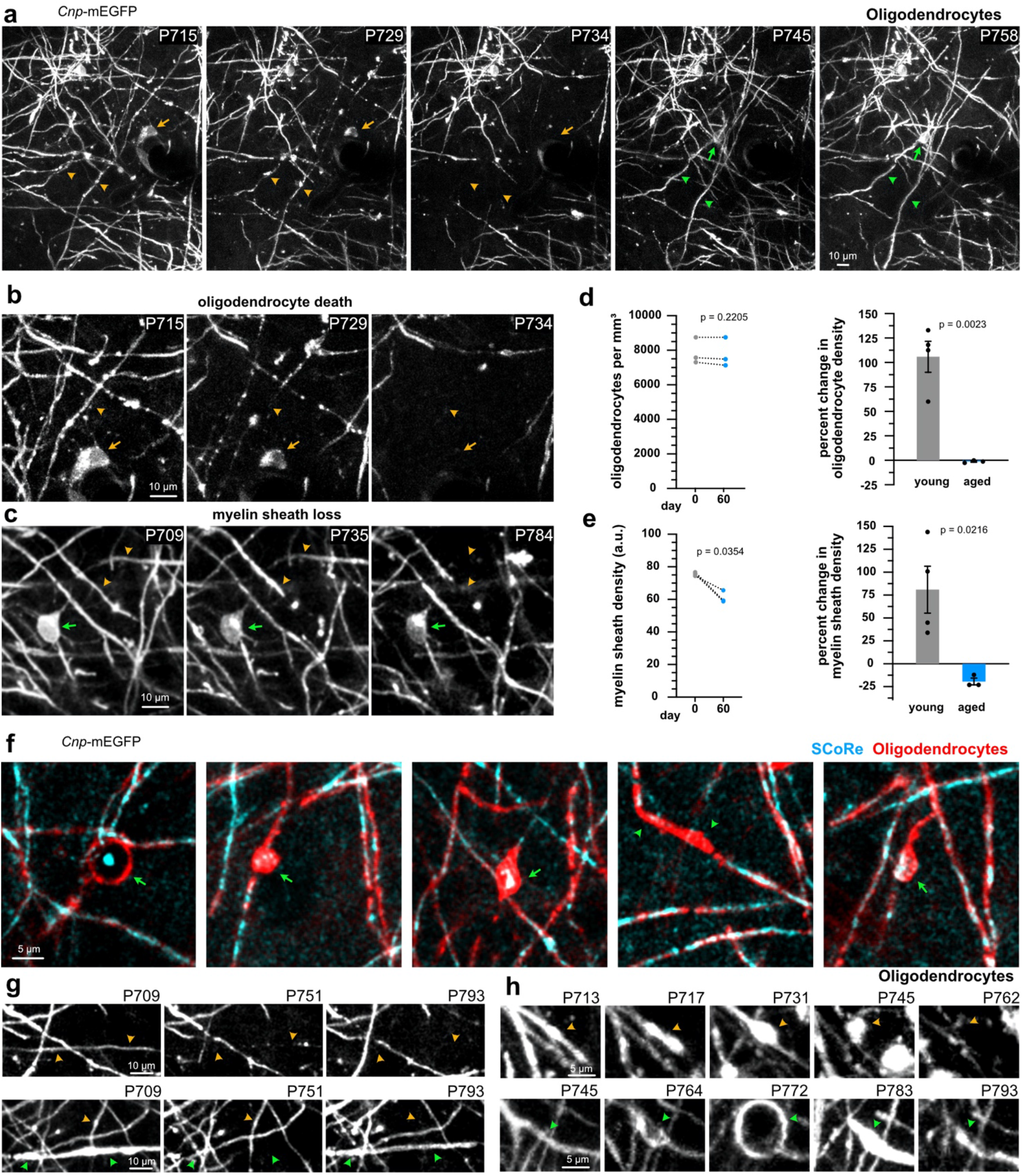
Age-related myelin degeneration occurs via sheath loss and cell death with limited repair. **a**) Oligodendrocyte death (yellow arrows) and myelin degeneration (yellow arrowheads) followed by the emergence of a newly generated oligodendrocyte (green arrows) and remyelinating sheaths (green arrowheads). **b**) Myelin pathology and degeneration coinciding (yellow arrowhead) with cell death (yellow arrow), cropped from the cell in panel (**a**) **c)**Myelin sheath degeneration without cell death (green arrow). **d**) Oligodendrocyte cell densities in aged mice over 60 days (n = 3 mice, paired t-test) and a comparison of this percent change in oligodendrocyte density between young and aged mice (young n = 4 mice, aged n = 3 mice, unpaired t test, error bars are SEM). **e**) Myelin sheath densities in aged mice over 60 days (n = 3 mice, paired t-test). Myelin sheath density increased in young mice between day 0 and day 60, while decreasing in aged mice (young n = 4 mice, aged n = 3 mice, unpaired t-test, error bars are SEM). **f**) Examples of commonly observed myelin pathology in aged mice including spheroid and myelin balloons (arrows) and loss of compaction (arrowheads), visualized by loss of SCoRe signal. **g**) Representative time series showing failed remyelination following myelin degeneration (orange arrowheads) and successful remyelination after sheath loss (green arrowheads). **h**) Severe myelin pathology, seen in aged mice, often resulted in full sheath degeneration (top, orange arrowheads), however sheath repair was also observed following spheroid formation (bottom, green arrowheads).

Myelin balloons are a common pathology that has been documented in *postmortem* aged brain tissue^37^. These were frequently observed across aged animals in our experiments and the presence of this type of structure often preceded full sheath degeneration (Fig. 7f-h,). In some cases, however, balloon formation would occur followed by what appeared to be repair of the damaged sheath without evidence of demyelination/remyelination (Fig 7h, Supplementary Fig. 6). These data suggested that the myelin sheath could potentially possess a mechanism for acute repair in response to the accumulation of fluid in the periaxonal space^38^ or between the layers of the myelin sheath.

Finally, to further determine the outcome of demyelination in the aging brain, we targeted single oligodendrocytes with 2Phatal. The reliability, timing of cell death, and morphological progression was identical to young animals (Fig 8 a-b, n = 36 young cells and 14 aged cells, Log-rank test for survival and unpaired t-test for days to cell death). However, unlike the young mice, we did not observe any remyelination or generation of new oligodendrocytes in response to 2Phatal induced demyelination (Fig. 8c). This resulted in a net loss of myelin sheaths around targeted cells in aged animals over the 60 days following 2Phatal photobleaching. While young animals were able to remyelinate due the generation of new oligodendrocytes over the same time frame (Fig. 8c-d, n = 10 young cells and 10 aged cells, two-way ANOVA with Sidak’s multiple comparisons test for sheath density and unpaired t-test for percent changes). Thus, after both spontaneous and experimentally induced demyelination in the aged brain, myelin repair is markedly absent due to the lack of new oligodendrocyte formation.

**Fig. 8.**
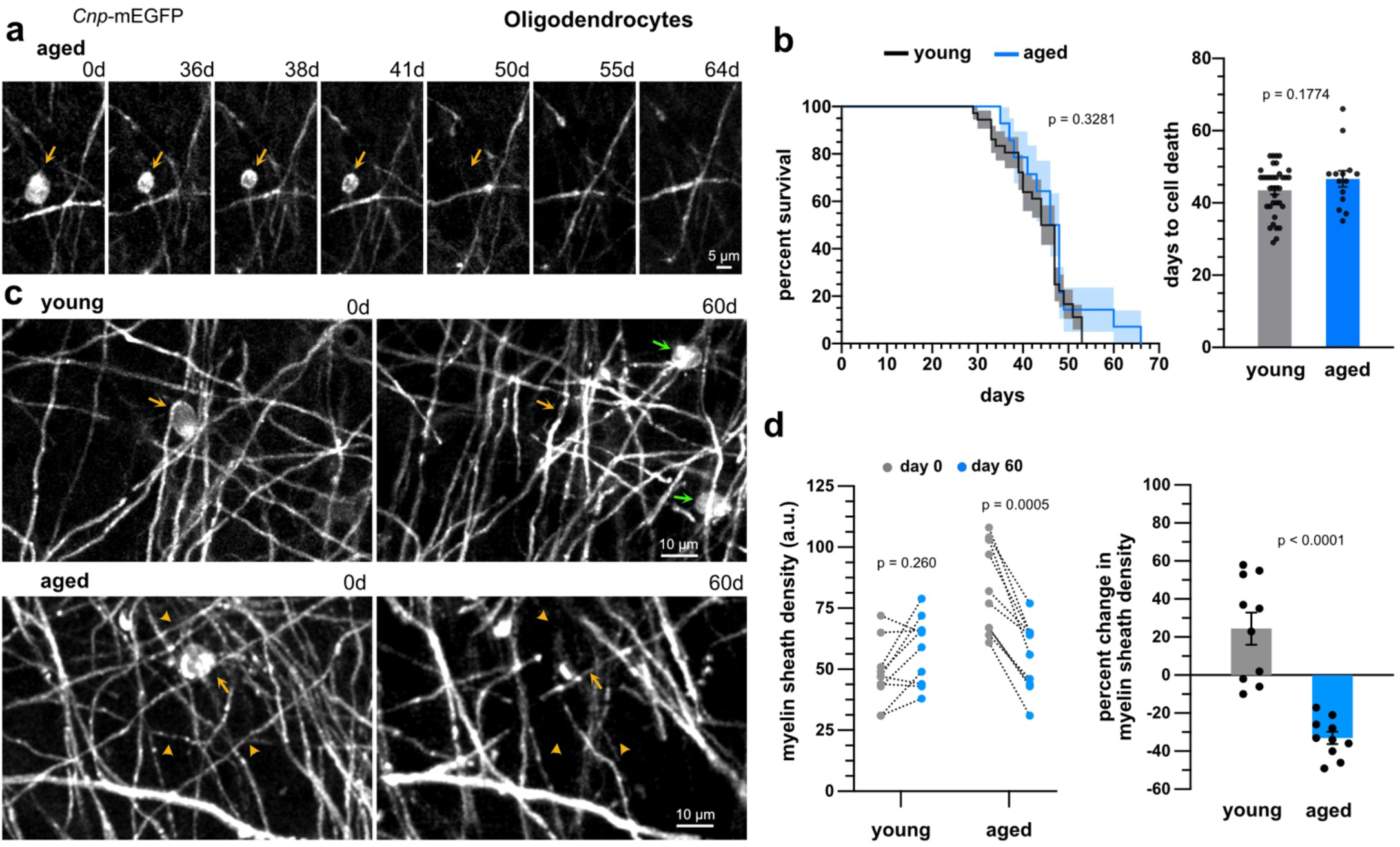
Failed myelin repair after oligodendrocyte 2Phatal in aging. **a**) *In-vivo* time series of an oligodendrocyte (arrow) degenerating following 2Phatal, in an aged mouse. **b**) Survival curves of oligodendrocytes after 2Phatal in young (n = 36 cells, 4 mice) and aged mice (n = 14 cells, 3 mice) (log rank (mantel cox) test, error bars are SEM). The average time to oligodendrocyte death, after 2Phatal, was not significantly different between young and aged mice (unpaired t test, error bars are SEM). **c**) Example images showing changes in myelin density (orange arrowheads) and new oligodendrocyte generation (green arrows) in the territory of an oligodendrocyte targeted with 2Phatal (orange arrows), in a young (top) and aged mouse (bottom). **d**) Quantification of the change in myelin sheath density, between day 0 and day 60, in the territory surrounding an oligodendrocyte targeted with 2Phatal, in young (n = 10 cells, 4 mice,) and aged mice (n = 10 cells, 3 mice, two-way ANOVA with Sidak correction for multiple comparisons). The change in myelin sheath density was significantly higher in young mice than aged mice (young n = 10 cells, 4 mice, aged n = 10 cells, 3 mice, unpaired t test, error bars are SEM).

## DISCUSSION

A major goal in myelin biology has been to understand the cellular and molecular mechanisms governing myelin repair. To this end, many demyelination models have been developed, including cytotoxic, immune, transgenic, and laser lesioning^3,13,29,39,40^. These approaches have all greatly added to our knowledge of cycles of demyelination and remyelination. However, many of these models illicit widespread systemic effects and some directly impact the ability of NG2-glia to differentiate and repair myelin until after the cessation of the insult^41,42^, a process which may or may not model disease or aging. In order to circumvent some of these challenges, we developed 2Phatal for use as a targeted, non-inflammatory, model of demyelination. Using this technique, combined with intravital optical imaging, we discovered several features of myelin degeneration and repair, including its effects on local NG2-glia behavior. First, using both fluorescence and label-free SCoRe microscopy, we discovered that structural myelin disruption, including loss of compaction, occurs days to weeks prior to full demyelination, indicating that normal axonal function is likely impaired earlier in myelin degeneration than previously thought. Second, we found that remyelination occurs more rapidly and efficiently along heavily myelinated axons, with some sheaths never fully losing compaction between degeneration and repair. Third, by analyzing the morphology of NG2-glia, we discovered that the majority of the myelin repair is carried out by a subset of highly branched NG2-glia. This represents the first evidence for local, functionally distinct, populations of NG2-glia in the context of remyelination. Finally, we found that age-related demyelination proceeds with similar spatiotemporal dynamics and morphological progression to both 2Phatal and cuprizone induced degeneration. Taken together, these data demonstrate *in-vivo* evidence of novel forms of rapid remyelination, carried out by a local subpopulation of NG2-glia. They further suggest that oligodendrocyte degeneration, in response to oxidative damage caused by 2Phatal, cuprizone, and aging, proceeds through conserved pathways, making 2Phatal a powerful new approach for investigating these mechanisms in the context of aging and disease.

The ability to determine functional changes in myelin over the course of degeneration is critical to understanding the acute consequences of myelin loss on neural circuitry. Temporal analyses of myelin compaction have been a challenge historically, as it has previously only been possible using electron microscopy in fixed tissue. Combining fluorescence and label-free SCoRe imaging, enables the direct characterization of sheath compaction over time, due to the specificity of SCoRe signal to areas of compact myelin. Evidence to support this specificity includes an absence of signal in non-compact regions, such as paranodes and proximal processes^33,35,36^, and little to no signal in the MBP-knockout shiverer mouse^35^, which lack the ability to produce compact myelin. Previous work has elegantly revealed the timing of sheath initiation by single oligodendrocytes^43^, molecular mechanisms controlling sheath compaction, and the state of compaction at different development states^44,45^. Building on this, we find that of myelin compaction lags behind sheath initiation by ~24 hours, in the context of remyelination, while complete compaction is not achieved until 3-5 days post initiation. Our use of SCoRe also uncovered a novel form of remyelination, with an almost uninterrupted transition between myelin sheath degeneration and formation of a new compact sheath. The lack of SCoRe loss notably decreased the time each axon spent without compact myelin. This rapid repair points to the existence of strong positive regulators of myelination produced by the axons on which it occurs. In fact, similar to propensity and timing for remyelination, myelin patterning was a strong predictor of remyelination in this way, as 94% of all sheaths exhibiting these dynamics were found on axons with partial or complete myelin patterns. In order for this rapid remyelination to occur, differentiating oligodendrocytes likely coopt the territory of degenerating sheaths prior to complete degeneration. Further investigation into the identity of the axons on which this process occurs will shed light on the factors that contribute to this form of remyelination

Previous work has focused primarily on understanding the molecular mechanisms that govern myelin loss, as well as how demyelination proceeds on a macro level. In the cuprizone model, for example, it is well established that demyelination occurs steadily over weeks, with peak myelin loss being achieved by around 6 weeks^42^. However, little is known about how demyelination proceeds on the single cell level, with recent intravital imaging experiments providing critical insight into the remyelination process after cuprizone cessation^27,28^. Our observations suggest that demyelination, in response to single cell oxidative DNA damage, occurs via the gradual loss of myelin sheaths from oligodendrocytes, rather than through single catastrophic death events. Adding to this finding, the steady loss of SCoRe signal over the course of degeneration, strongly suggests that decompaction occurs in advance of sheath loss, likely negatively impacting the function of ensheathed axons prior to demyelination. While there is evidence for decompaction in the aged brain^37^, the dynamics and functional consequences to the single axon are yet to be determined.

Recent studies have provided evidence for remyelination by mature oligodendrocytes, raising the question of whether or not oligodendrogenesis is required for remyelination to occur. This data comes from postmortem analysis of shadow plaques in MS tissue^46,47^ and from intravital imaging studies in mice exposed to cuprizone while undergoing a motor learning task^27^. Using our dual-reporter system, we found no evidence of pre-established myelinating oligodendrocytes participating in remyelination. Clearly the cellular scale and molecular insult of single cell 2Phatal demyelination is different from both cuprizone intoxication and MS which could account for this difference. Moreover, 100% of oligodendrocytes exposed to 2Phatal photobleaching end up dying, while damaged but surviving oligodendrocytes are likely to be the cells that are involved in the remyelination, at least in the context of cuprizone intoxication^27^. Future experiments investigating whether titration of 2Phatal photobleaching can result in cellular DNA damage and repair without resulting in cell death would provide a means to test this question. Furthermore, the extent to which motor learning induces remyelination by pre-established oligodendrocytes could also be tested in the context of 2Phatal demyelination.

Genetically distinct populations of NG2-glia are known to exist regionally and temporally. For example, studies using RNA-seq and electrophysiology have demonstrated differential expression of ion channels, in NG2-glia, between grey and white matter and at different developmental stages^15^. Although diversity, such as this, does appear to influence NG2-glia behavior regionally, little is known about functional heterogeneity within local populations. Our findings indicate that myelin repair is largely carried out by a subset of highly branched NG2-glia. NG2-glia with simpler morphology were less likely to participate in remyelination. As our imaging was limited to layer I of the somatosensory cortex and highly branched cells were found adjacent to cells with simpler morphology, this suggests that functional heterogeneity can exist in a spatially confined region of the mammalian brain. Furthermore, as morphological complexity was predictive of cell behavior weeks after analysis, it seems unlikely that our observations were due to transient morphological changes resulting from active differentiation or cell death. Consistent with our findings, a recent report also correlated NG2-glia morphology with cellular function in zebrafish^17^. Whether or not these observations are the result of differences in cellular microenvironment, signaling between neurons and subsets of NG2-glia, or represent local genetic diversity within the NG2-glia population remains unclear.

Lastly, myelin degeneration is known to occur in the aging brain, likely contributing to cognitive decline^37,48^. Oligodendrocyte loss coupled with failed differentiation of NG2-glia leads to a steady decline of myelin^49,50^. This results in the accumulation of debris, made worse by decreased phagocytic activity and failed degradation of lipids by resident glia^33,51,52^. While the features of age-related myelin loss have been extensively characterized in postmortem tissue from rodents and primates, the cellular dynamics, molecular mechanisms involved, and its functional impacts on the CNS remain largely unanswered^3,5,48^. Our analysis of myelin degeneration, in the context of 2Phatal, cuprizone and aging, revealed remarkable similarities in both the spatiotemporal dynamics and pathology across these conditions. Intriguingly, oxidative stress and DNA damage are thought to play a role in oligodendrocyte death in both cuprizone and aging^2,3,48,53^. Given the nature of 2Phatal, it is likely that direct oxidative DNA damage is responsible for its induction of cell death. Taken together, these data suggest that the drawn-out mechanism of oligodendrocyte degeneration we have described represents a conserved, intrinsic response to oxidative damage by myelinating oligodendrocytes. This, combined with its versatility, makes 2Phatal an ideal method for investigating the precise cellular and molecular mechanisms involved in oligodendrocyte degeneration in the aged brain.

## ACKNOWLEDGEMENTS

This work was supported by the following grants from the National Institutes of Health: R00-NS099469 and P20-GM113132 and by a New Vision Award through the Donors Cure Foundation and a Fay/Frank Seed Grant from the Brain Research Foundation to R.A.H. We thank Dartmouth colleagues Leeza Petrov and Xhoela Bame for contributions to data analysis. We thank members of the Hoppa Lab at Dartmouth for critical feedback throughout the project.

## AUTHOR CONTRIBUTIONS

T.W.C and R.A.H conceived of, designed, and performed all experiments and the majority of data analysis and quantification. G.E.O. performed NG2-glia morphological and migration analyses and contributed to quantification associated with sheath and node remyelination. E.P. contributed to NG2-glia fate analysis and imaging data collation. T.W.C and R.A.H. wrote the paper and R.A.H. directed the study.

## METHODS

### Animals

All animal procedures were approved by the institutional animal care and use committee (IACUC) at Dartmouth College. The following mouse strains were purchased from Jackson labs and crossed to generate the double and triple transgenic mice used in this study: *Cnp*-mEGFP^32^ (JAX #026105), *Cspg4*-creER^54^ (JAX #008538), *Cx3cr1*-creER^55^ (JAX #020940), floxed tdTomato Ai14^56^ (JAX #007914). Mice were housed in a 12/12 light/dark cycle in a temperature and humidity-controlled animal vivarium with food and water provided ad libitum. For all experiments (except aging) mice were 6-8 weeks old at the start of the experiments with control mice being aged matched. Aged mice were defined as 22-24 months old. Both male and female mice were used in this study.

### Surgical procedures

All *in vivo* imaging was done using chronic cranial window preparations. In short, animals were anesthetized by intraperitoneal injection of ketamine (100 mg/kg) and xylazine (10 mg/kg). The skin covering the skull was shaved and sterilized before removal. A high-speed drill was used to perform a 3-4 mm craniotomy and the skull was replaced by a #0 cover glass. For 2Phatal experiments, Hoechst 33342 nuclear dye (ThermoFisher #H3570) was topically applied to the pial surface, before placing the cover glass. A nut was attached to the skull with cyanoacrylate glue and embedded in dental cement to enable repeated animal immobilization for imaging. Mice were given carprofen (50 mg/kg), subcutaneously, following the surgery and at 24- and 48-hours.

### 2Phatal

For targeted two-photon apoptotic targeted ablations (2Phatal), cranial windows were performed as described above. Hoechst 33342 was topically applied (0.1 mg ml^−1^ diluted in PBS) to the pial surface for 5 minutes before the cover glass was secured. Control mice received identical dye applications without subsequent 2Phatal. Mice were allowed to recover for 24 hours to ensure proper nuclear labeling. Each position was imaged sequentially on a two-photon microscope using 775nm, 920nm and 1040nm laser wavelengths and the channels were overlayed, to identify mature oligodendrocyte nuclei. To induce 2Phatal, a single ROI (8×8 μm^2^) was drawn around the identified nucleus. The laser wavelength was set at 775nm, with a dwell time of 100 μs. For each ablation, we employed the same time-series parameters of 125 scans, lasting in total 3.72 seconds. Following 2Phatal, each location was imaged again using 775nm and 920nm to visualize photobleaching and ensure no membrane disruption or laser burning was caused during each photobleaching event. To quantify photobleaching, the average fluorescence intensity was calculated for each time point. These values were then normalized to the average intensity of the first scan to get a percent change in fluorescence intensity over time.

### Cuprizone demyelination

Cranial window preparations were performed on *Cnp*-mEGFP mice before a recovery period of 3 weeks. Following this, baseline, day 0, images were taken, and the mice were placed on a 0.2% (w/w) cuprizone (Sigma Aldrich #C9012) diet mixed with ground chow for 6 weeks. Fresh food was provided every 2-3 days. After 6 weeks of treatment, mice were placed back on normal, control, powdered chow for 4 weeks. Combined fluorescence and SCoRe *in vivo* imaging were conducted weekly using an upright laser scanning confocal microscope (Leica SP8) with a 20x water immersion objective (Leica NA 1.0).

### Imaging

All fluorescence *in vivo* imaging was performed using an upright laser scanning confocal microscope (Leica SP8) with a 20x water immersion objective (Leica NA 1.0) or a two-photon microscope (Bruker) equipped with an Insight X3 femtosecond pulsed laser (Spectra Physics) and a 20x water immersion objective (Zeiss NA 1.0). Spectral confocal reflectance (SCoRe) microscopy was done using the upright laser scanning confocal microscope. Reflection signal was gathered from 448nm, 488nm, 552nm, and 637nm lasers and overlayed to identify compact myelin. For confocal fluorescence microscopy, 488nm was used to excite mEGFP and 552nm was used to excite tdTomato. Fluorescence and SCoRe images were acquired sequentially. For two-photon fluorescence microscopy, 775nm was used to excite Hoechst nuclear dye, 920nm was used to excite mEGFP, and 1040nm was used to excite tdTomato. Z-stacks were taken with a step size of 1.5μm. Mice were imaged as described in the text and figures.

### Microglia response to 2Phatal

To investigate the acute response of microglia to 2Phatal photobleaching, targeted and control non targeted adjacent oligodendrocytes were selected. Cropped max projection images were analyzed and the fluorescence intensity was measured of the microglia signal in a 50um radius region of interest centered on the targeted or control oligodendrocyte before and 20 minutes after photobleaching as previously described^30^. Changes in microglia fluorescence were calculated and compared between the control non-bleached and 2Phatal cells using unpaired t-tests.

### NG2-glia fate

To quantify NG2-glia fate in *CNP*-mEGFP:*Cspg4*-creER:tdTomato mice, HyperStacks were created for days 28-60 after 2Phatal or in similarly aged control mice that did not have any oligodendrocytes targeted with 2Phatal. Z projection images were created and registered, and cell behavior and the fate were analyzed for each cell with NG2-glia morphology. Potential fates noted were 1) remained NG2-glia 2) cell death and 3) oligodendrocyte differentiation as evidenced by expression of *CNP*-mEGFP. Cell division events were also noted for all analyzed cells. Lineage diagrams were constructed for each cell (examples in Fig. 3e) and differences in the fate of NG2-glia in control mice and mice that had oligodendrocytes targeted via 2Phatal were determined using unpaired t-tests as indicated in the text and figures. For analysis of NG2-glia fate adjacent to or away from dying oligodendrocytes, a region of interest (ROI), 150μm in diameter, was drawn around targeted oligodendrocyte cell soma. NG2-glia were classified as away from or adjacent to targeted cells if they were outside or inside of these ROIs on day 28 after 2Phatal.

### NG2-glia migration

For analyses of NG2-glia migration in control and 2Phatal conditions, images were aligned for the whole time series of day 28 to day 60 and made into a HyperStack. The image stack was then opened in Fiji using the TrackMate plugin. Calibration settings were then inputted to convert pixels to microns. The LoG detector was then selected and an estimated blob diameter of 13μm was used for the cell body and the sub pixel localization was turned off. Once TrackMate detected the “blobs” in the image stack, an automatic threshold was applied. The detected blobs are overlayed onto the image using the HyperStack displayer. The Simple LAP Tracker was then selected to track the movement of the cells throughout the time series. Initially the Linking Max distance was set to 15μm, Gap-closing max distance was set to 15μm, and the Gap-closing max frame gap was set to 2. The tracks will then be overlayed onto the HyperStack displayer. If breaks in the track for individual cells were detected after visual inspection the parameters in the Simple LAP Tracker were adjusted. After all the tracks were confirmed to follow the movement of the cell, TrackMate was used determined the total distance traveled for each cell. These values were then compiled and analyzed in relation to condition and cell fate.

### NG2-glia morphology

Sholl analyses were conducted on max projection cropped images using the Sholl plugin on ImageJ. Before analysis all images were converted to an 8-bit format with a grey overlay, smoothed, and auto threshold. A straight line was drawn from the center of the cell to the outer most part of the image. The starting radius was set to 2μm and the end radius was set to 100μm. 2μm increments were used for analysis. In some cases, single cells were traced using simple neurite tracer plugin in Fiji and total process length in microns was determined from these cell reconstructions.

### Oligodendrocyte death and myelin sheath dynamics

To facilitate analysis, time series of each position were combined into HyperStacks using Fiji. For the quantification of survival curves in the 2Phatal and cuprizone models (Fig. 1h,i), cell death was defined as the first time point in which there was no longer evidence of the cell soma as seen by mEGFP fluorescence. To quantify the survival curve and average time to cell death for 2Phatal, every targeted cell was tracked from day 0 to degeneration of the cell soma. For cuprizone, every oligodendrocyte in the field of few on day 0 was counted and evaluated at subsequent time points for degeneration. However, only cells that died over the course of the experiment were used to calculate average time to cell death. For the quantification of changes in oligodendrocyte soma area (Fig. 1g), ROIs were drawn around each cell in Fiji, using the Selection Brush Tool, and used to calculate soma area at timepoints 1-3 days apart. In cases where the soma was in focus across multiple z slices, the slice with the largest area was used. Cells were excluded from soma area analysis if there were timepoints where the local image quality of individual cells was insufficient to acquire an accurate area measurement.

For the quantification of sheath loss of individual oligodendrocytes (Fig. 2b), the proximal processes of targeted oligodendrocytes were traced from the cell soma to the adjoining internodes. Sheaths were excluded if there was any doubt as to the cell of origin. As with cell death, each sheath was marked as degenerated on the first timepoint with no evidence of the sheath seen by mEGFP fluorescence. For the quantification of changes in sheath length and SCoRe:GFP ratio of degenerating cells (Fig. 2), a subset of the previously identified sheaths was analyzed. The Segmented Line Tool was used in Fiji to trace each sheath. For SCoRe measurements, each reflection wavelength was combined into a single channel, used to calculate lengths by summing the segments of compact myelin. Breaks in SCoRe signal, ≥1.5μm, were defined as sections of uncompact myelin which were not added to the SCoRe length sum of each sheath. SCoRe:GFP ratio was calculated by dividing the sum of SCoRe measurements by the measured length of each sheath (mEGFP channel). In depth quantification of remyelination (Fig. 6d) was done in the same manner.

For the quantification of degeneration and remyelination across each position (Fig. 5c), mEGFP+/tdTomato− sheaths were blindly selected from the first time point), independent of the cell of origin, and evaluated on day 60 for stability (mEGFP+/tdTomato−), remyelination (mEGFP+/tdTomato+) or degeneration without repair (lost). For the quantification of node stability (Fig. 5j,k), nodes of Ranvier were blindly selected (both hemi nodes were mEGFP+/tdTomato−), independent of the cells of origin, and evaluated on day 60 as before.

For the quantification of remyelination efficiency and time to remyelination, sheaths, originating from targeted cells, were tracked over the time series to determine when degeneration took place and when/if they were remyelinated (Fig. 5e,f). Patterning was determined on the first time point, based on the presence of adjacent internodes. To quantify time to remyelination, we calculated the time between disappearance of each sheath and initiation of repair, using mEGFP and tdTomato fluorescence. To quantify the prevalence of each identified method of remyelination (compaction and GFP lost; compaction lost, GFP maintained; compaction and GFP maintained), we tracked each sheath through every time point (identified above) and documented mEGFP and SCoRe loss during remyelination.

### Oligodendrocyte and myelin sheath density in young and aged animals

Oligodendrocyte density was determined by counting the total number of oligodendrocyte soma in at least 2 separate images from 4 young mice and 3 aged mice at two time points 60 days apart. Image dimensions were used to calculate the density of oligodendrocytes in each animal in cells / mm^3^. For measurements of myelin sheath density in young and aged control and 2Phatal targeted cells 100μm × 100μm images centered on a single oligodendrocyte soma were created and mean fluorescence intensity measurements were recorded from auto-thresholded images.

## COMPETING INTERESTS

The authors declare no competing interests

## DATA AVAILABILITY

All relevant data is available from the authors upon reasonable request.

## Notes

### Competing Interest Statement

The authors have declared no competing interest.

