## Supplementary Information for "Age and axon-specific forms of cortical remyelination by divergent populations of NG2-glia"

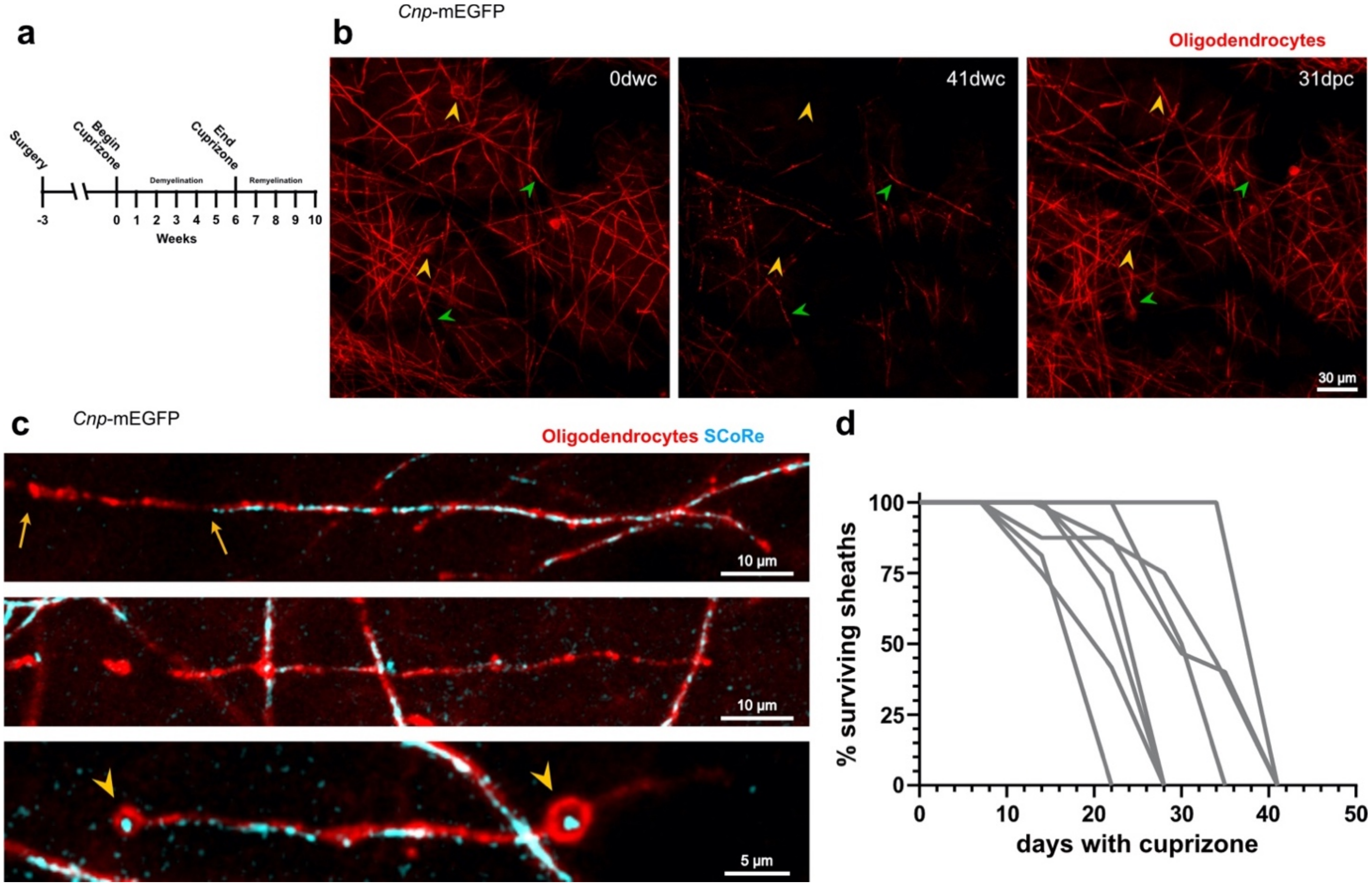


**Supplementary Figure 1: Cuprizone-induced myelin pathology and dynamics of degeneration**

**a)** Paradigm used for cuprizone induced demyelination experiments. Mice were fed 0.2% w/w cuprizone mixed in ground chow. *In-vivo* imaging was done weekly, starting at week 0, through week 10. **b)** Cuprizone induced widespread demyelination and oligodendrocyte cell loss (orange arrowheads) after 41 dwc (days with cuprizone). Remyelination was largely complete by 31 dpc (days post cuprizone). Green arrowheads denote points of reference for position orientation. **c)** Representative images of common myelin pathology observed during cuprizone intoxication, including partial loss of compaction (top, between arrows), visualized by a lack of SCoRe signal, complete loss of compaction with myelin debris (middle), and spheroid formation (bottom, arrowheads). **d)** Temporal dynamics of sheath degeneration during cuprizone intoxication. Each trace represents sheaths produced from a single oligodendrocyte (n = 8 cells, 116 sheaths, 3 mice).

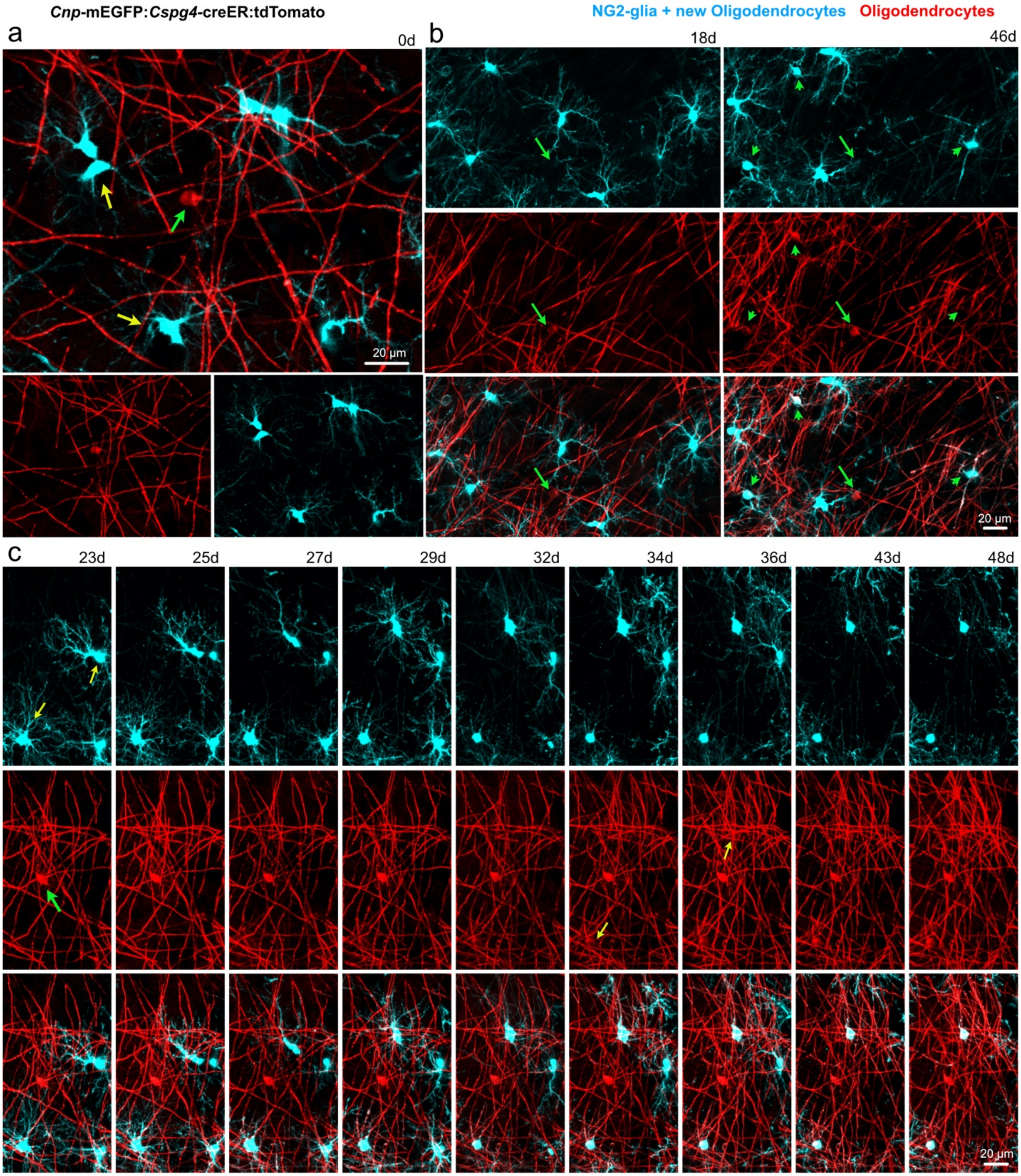


**Supplementary Figure 2: The *Cnp*-mEGFP:*Cspg4*-creER:tdTomato mouse line identifies newly generated oligodendrocytes.**

**a)** Example *in-vivo* image showing a previously generated, tdTomato-, oligodendrocyte (green arrow) and tdTomato+ NG2-glia (yellow arrows) at day 0. **b)** Time series showing the persistence of a previously established, mature, oligodendrocyte (arrow) and the generation of new dual labeled mEGFP+tdTomato+ oligodendrocytes (arrowheads) between day 18 and day 46. **c)** Time series showing the differentiation of new dual labeled oligodendrocyte (yellow arrows) over the course of the experiment, while a previously established single labeled oligodendrocyte (green arrow) is maintained.

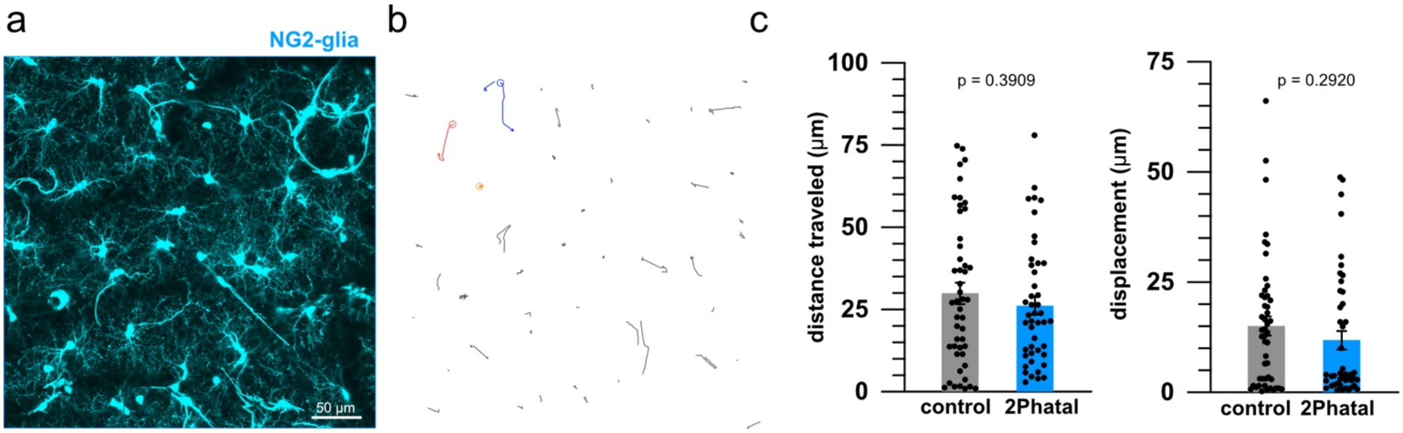


**Supplementary Figure 3: Local NG2-glia migration is unaffected by 2Phatal induced cell death of oligodendrocytes.**

**a)** Representative MAX projection of a single time point, showing NG2-glia (cyan), used to determine the migration of NG2-glia over time. **b)** Example NG2-glia migration tracks between day 28 and day 60 from a single imaging location. Each line represents the migration track of a single cell. **c)** The total distance traveled (left) and net displacement (right) of NG2-glia in control mice compared to mice targeted with 2Phatal. There was no significant difference in overall migration behavior between the two groups (control n = 48 cells, 2Phatal n = 45 cells, unpaired t test, dots indicate single cells).

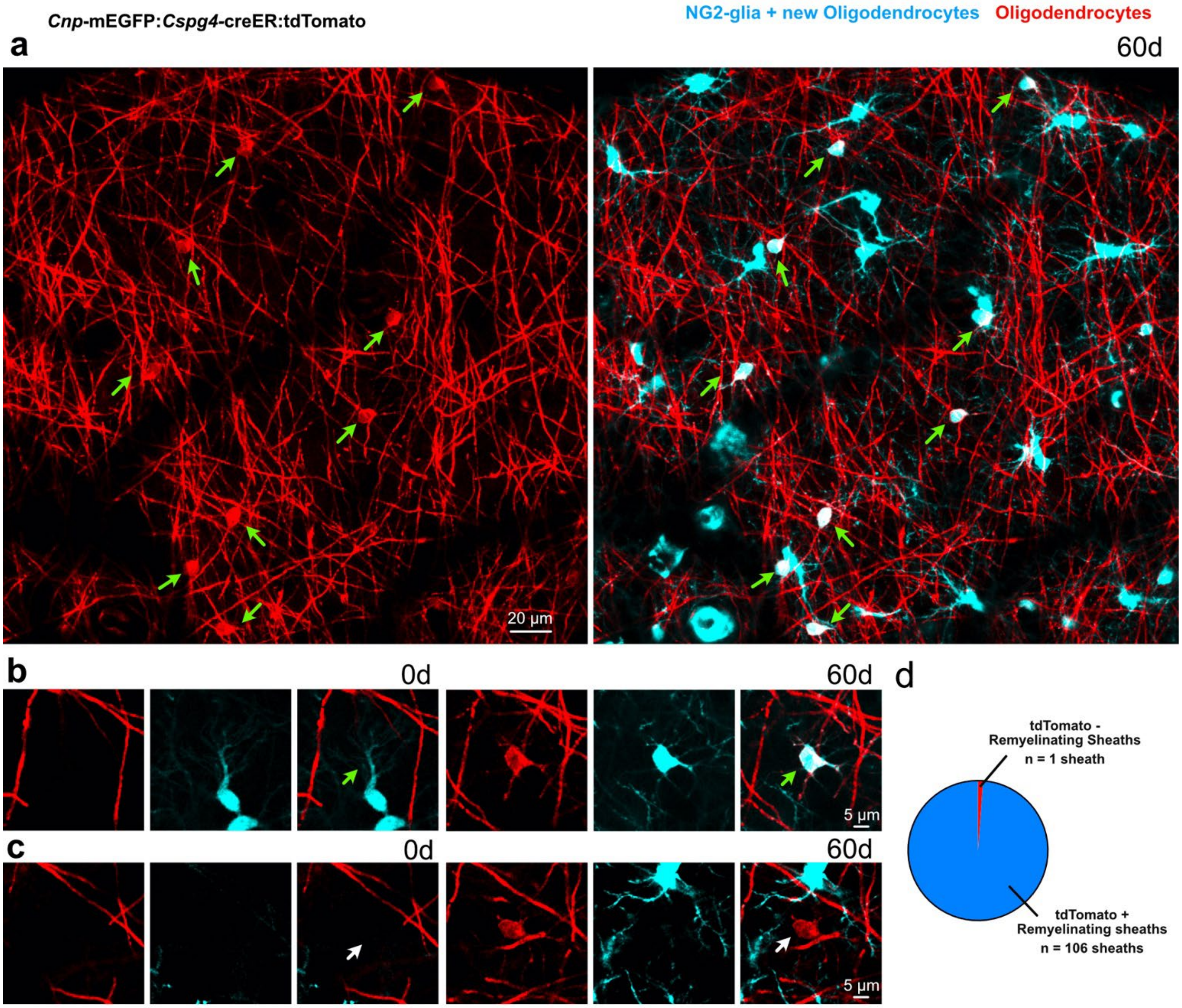


**Supplementary Figure 4: Efficient Cre recombination allows fate mapping of oligodendrocyte generation in vivo.**

**a)** Representative image of a *Cnp*-mEGFP:*Cspg4*-creER:tdTomato mouse at day 60 showing all surviving oligodendrocytes (arrows) are dual labeled with mEGFP+ and tdTomato+. For this example all cells at day 0 that were mEGFP+ were targeted with 2Phatal **b)** Time series showing the presence of a new mEGFP+/tdTomato+ oligodendrocyte (green arrow) at day 60. **c)** During all experiments we encountered only a single tdTomato- oligodendrocyte, that was generated between day 0 and day 60 (white arrow). **d)** Of the 107 remyelinating sheaths analyzed, a single sheath was tdTomato-. As it was in the vicinity of the tdTomato- cell shown in **c**, it is likely to have originated from that cell.

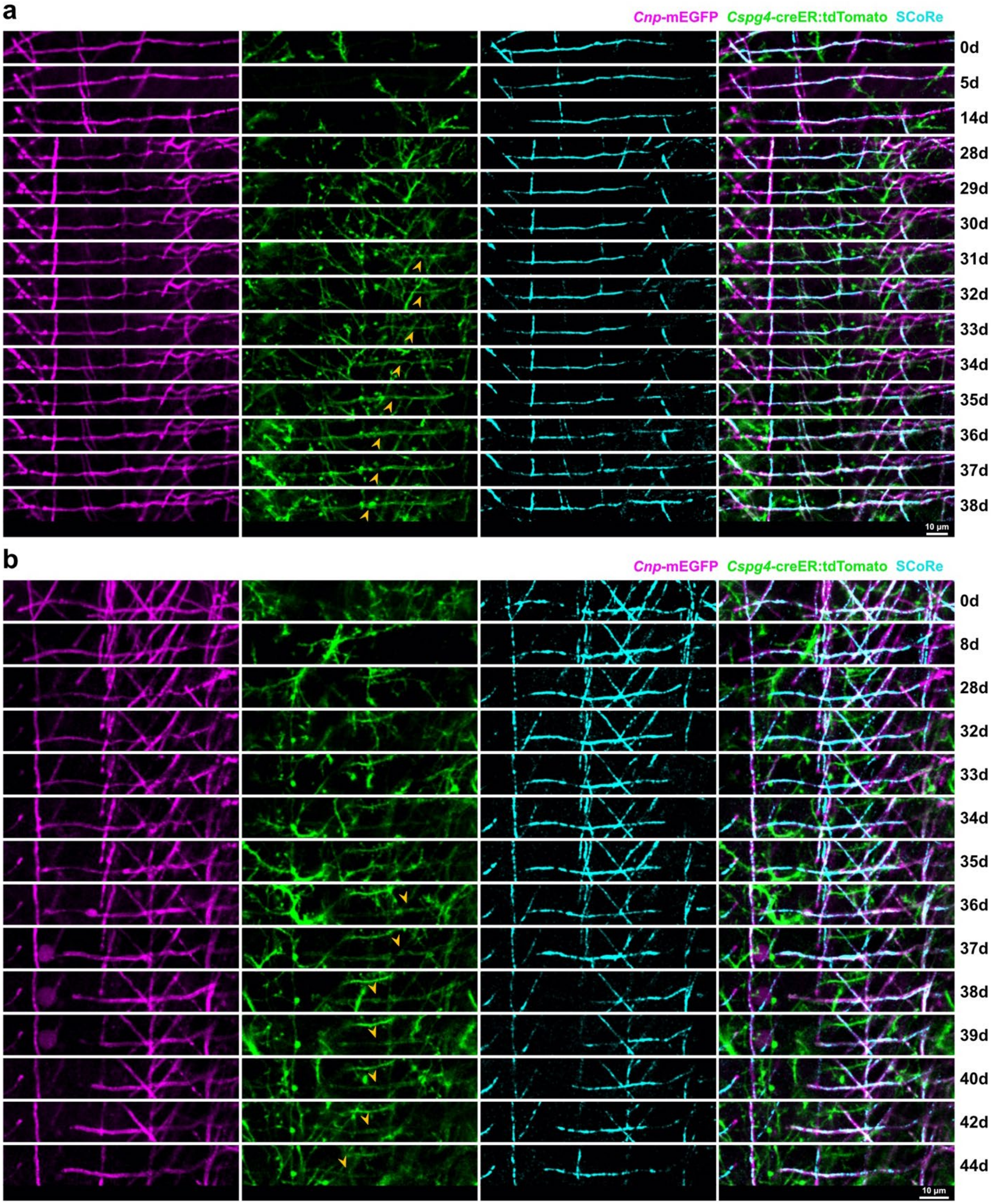


**Supplementary Figure 5: Example time series of remyelination without SCoRe loss**
**a)** Original images used to generate the traced time series shown in Fig. 6c. The initiation of remyelination is evident by emergence of tdTomato signal (arrowheads) on day 31. **b)** Additional example of remyelination without score loss (not shown in the main text).

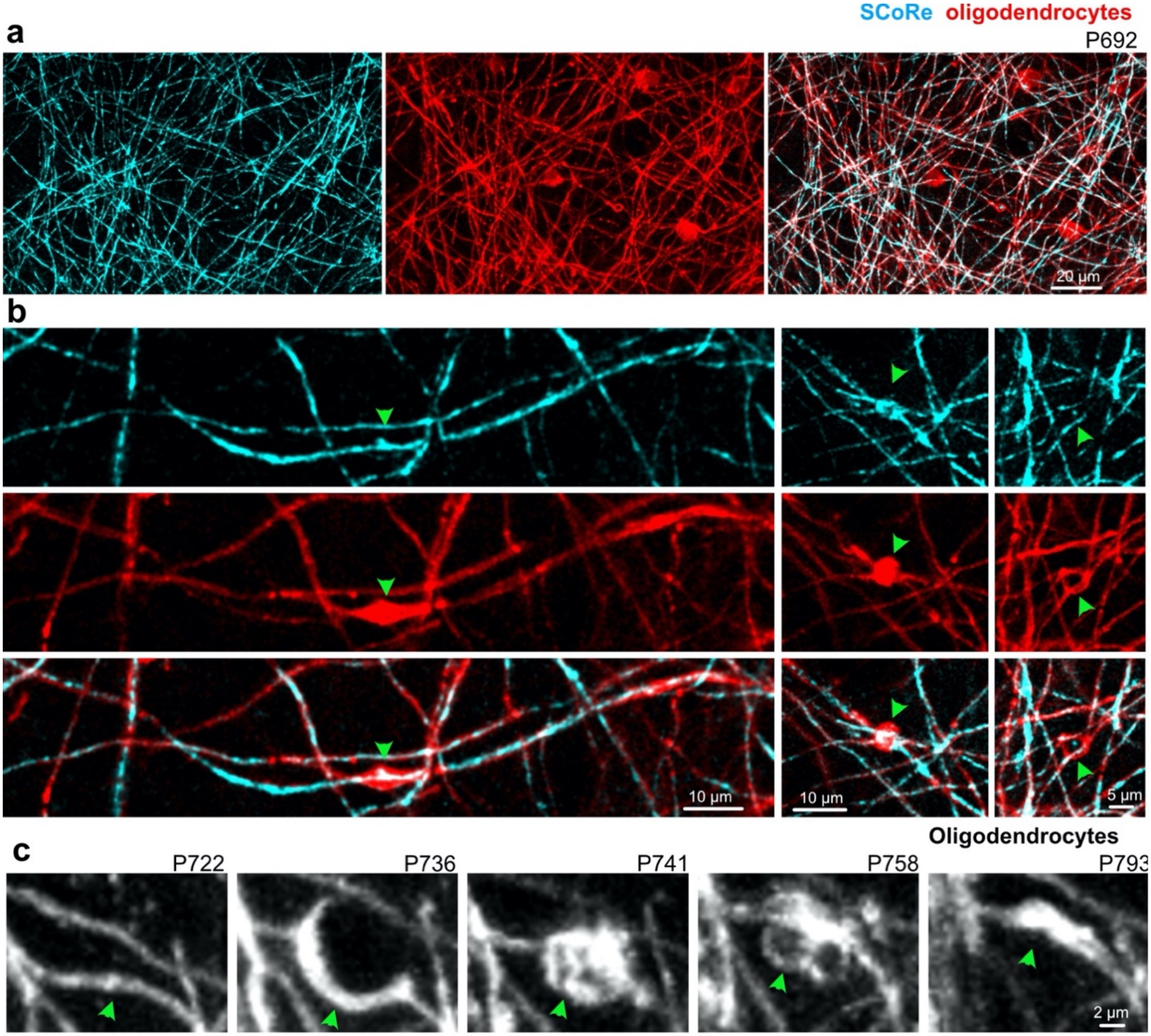


**Supplementary Figure 6: Common myelin pathology observed in aged mice**

**a)** *In-vivo* image of myelin, in layer I of the somatosensory cortex, in an aged mouse, acquired using SCoRe and fluorescence microscopy. **b)** Aged mice displayed widespread myelin pathology (arrowheads) including myelin swellings (left), debris accumulation (middle), and spheroids (right). **c)** Timeseries showing spontaneous spheroid formation and spontaneous repair (arrowhead).
